# Adaptation to spindle assembly checkpoint inhibition through the selection of specific aneuploidies

**DOI:** 10.1101/2022.10.04.510607

**Authors:** Manuel Alonso Y Adell, Tamara C. Klockner, Rudolf Höfler, Lea Wallner, Julia Schmid, Ana Markovic, Anastasiia Martyniak, Christopher S. Campbell

**Affiliations:** Department of Chromosome Biology, Max Perutz Labs, University of Vienna, Vienna Biocenter (VBC), A-1030 Vienna, Austria

**Keywords:** Key words: Aneuploidy Patterns – Chromosomal Instability – Spindle Assembly Checkpoint – Drug Resistance Mechanisms – MPS1

## Abstract

Both the presence of an abnormal complement of chromosomes (aneuploidy) and an increased frequency of chromosome missegregation (chromosomal instability) are hallmarks of cancer. Analyses of cancer genome data have identified certain aneuploidy patterns in tumors; however, the bases behind their selection are largely unexplored. By establishing time-resolved long-term adaptation protocols, we found that human cells adapt to persistent spindle assembly checkpoint (SAC) inhibition by acquiring specific chromosome arm gains and losses. Independently adapted populations converge on complex karyotypes, which over time are refined to contain ever smaller chromosomal changes. Of note, the frequencies of chromosome arm gains in adapted cells correlate very well with those detected in cancers, suggesting that our cellular adaptation approach recapitulates selective traits that dictate pan-cancer aneuploidy patterns. We further engineered specific aneuploidies to determine the genetic basis behind the observed karyotype patterns. These experiments demonstrated that the adapted and engineered aneuploid cell lines limit CIN by extending mitotic duration. Heterozygous deletions of key SAC and APC/C genes recapitulated the rescue phenotypes of the monosomic chromosomes. We conclude that aneuploidy-induced gene dosage imbalances of individual mitotic regulators are sufficient for altering mitotic timing to reduce CIN.

## Introduction

The faithful segregation of replicated genetic material to daughter cells is a fundamental requirement of all living beings. Erroneous chromosome segregation during mitosis results in gains and losses of chromosomes and chromosome arms, resulting in aneuploidy. An increased rate of such chromosome missegregation events is termed chromosomal instability (CIN). Aneuploidy is associated with growth abnormalities and inviability in many organisms (reviewed in Torres, Williams, and Amon 2008; Williams and Amon 2009). In humans, all autosomal chromosome losses are embryonic lethal and only a few chromosome gains, such as trisomy 21, are compatible with life (Hassold and Hunt 2001). On the cellular level, aneuploidy results in dosage changes of hundreds of genes at once, leading to a wide variety of phenotypes including proteotoxic stress, genome instability and cell cycle arrest (reviewed in Zhu et al. 2018; Chunduri and Storchová 2019).

Despite these adverse effects, aneuploidy is found in ∼90 % of solid tumors and ∼70 % of hematopoietic cancers, making it one of the most common types of genetic alterations in cancer (Weaver and Cleveland 2006). Aneuploidy and CIN are associated with tumor progression and metastasis formation (Carter et al. 2006; McGranahan et al. 2012; Shukla et al. 2020). Experiments from yeast to human cells have shown that certain aneuploidies can be advantageous under specific stress conditions (Rutledge et al. 2016; Ravichandran et al. 2018; Salgueiro et al. 2020; Selmecki, Forche, and Berman 2006). Recent studies demonstrated that temporary induction of CIN confers resistance to chemotherapeutic drugs (Ippolito et al. 2021; Lukow et al. 2021). However, the selective forces underlying the retention of specific aneuploidies are currently not well understood.

Cancers have characteristic aneuploidy patterns. For example, certain aneuploidies like the gain of chromosome arms 8q and 20q are highly frequent across many different cancer types (Beroukhim et al. 2010). Other aneuploidies are only prevalent in specific cancer types, such as chromosome arm 3p loss in squamous cancers or 13q gain in gastrointestinal tumors (Taylor et al. 2018). The complexity of cancer karyotypes makes it difficult to determine the selective advantage of specific aneuploidies. However, many studies have shown interesting correlations in aneuploidy patterns. For example, a seminal study indicated that the distribution of tumor suppressors and oncogenes across chromosomes can predict whether a particular chromosome is more likely to be gained or lost (Davoli et al. 2013). Recent breakthroughs in experimental models for Ewing sarcoma in mice and human cells have found a partial contribution in the Rad21 and Myc genes for the selection of chromosome 8 trisomy (Su et al. 2021). However, there are currently very few bottom-up methods to identify the adaptive advantage of specific aneuploidies in human cells and identify the underlying responsible genes.

The sources of CIN in cancer have long been elusive. Although many potential sources of CIN have been identified, such as supernumerary centrosomes, whole genome duplication, and replication stress, the degree to which each of these abnormalities contribute to chromosome missegregation is still currently unclear (reviewed in Sansregret, Vanhaesebroeck, and Swanton 2018). What is well established is that chromosome segregation errors are largely prevented by the spindle assembly checkpoint (SAC). The SAC is a surveillance mechanism monitoring accurate and timely attachment of chromosomes to the mitotic spindle (reviewed in Musacchio and Salmon 2007; Lara-Gonzalez, Pines, and Desai 2021; Silva et al. 2011; Pachis and Kops 2018). SAC activation inhibits the anaphase promoting complex/cyclosome (APC/C), preventing cells from entering anaphase until all of the chromosomes are attached to spindle microtubules (Watson et al. 2019). Disruption of SAC activity can lead to tumorigenesis in mice, demonstrating the importance of the SAC as a tumor suppression mechanism (Sotillo et al. 2007). Interestingly, a number of oncogenic viruses downregulate the activity of SAC components (Jin, Spencer, and Jeang 1998; Sun et al. 2014; Shirnekhi et al. 2017). However, it is currently unclear if and how persistent SAC downregulation influences complex aneuploid karyotype formation in human cells.

In this work, we used a time-resolved adaptation assay based on long-term inhibition of the SAC kinase MPS1 to analyze how complex aneuploid karyotypes change over time in response to CIN. We found that complex karyotypes converge on very similar aneuploidy patterns. These complex karyotypes are then refined towards smaller copy number alterations (CNAs). Intriguingly, the frequencies of chromosome gains in the adapted cells correlate with those seen in cancer, suggesting general advantages that are largely independent of the cell type and cellular environment. We then used CRISPR/Cas9-based engineering of monosomic chromosomes to determine the genetic bases behind frequently acquired aneuploidies. We identified specific monosomies that directly rescue SAC inhibition. We show that these monosomies increase the mitotic duration, and that changing the dosage of single genes is sufficient to reduce CIN. Together, these results demonstrate that alterations in single genes on aneuploid chromosomes are sufficient to affect mitotic timing and suppress SAC inhibition.

## Results

### Long-term MPS1 inhibition leads to selection of specific aneuploidies and karyotype refinement over time

To study how human cell karyotypes change over time in the presence of high rates of chromosome missegregation, we continuously treated p53-deleted cell lines with the MPS1 inhibitor reversine over 30, 60 and 90 days (Fig. 1a). Reversine was added at concentrations high enough to act simultaneously as a CIN inducing agent and a strong selective pressure (Extended Data Fig. 1a,b). Before initiating the adaptation process, we first engineered human cell lines suitable for observing adaptation to high rates of CIN. We started with six near-diploid human cell lines that comprise three categories: myeloid leukemia (diploidized HAP1, hereafter called dipHAP1, and EEB), colorectal cancer (HCT116 and DLD1) and epithelial non-cancer (hTERT-RPE1 and hTERT-HME1). Near-diploid cell lines were chosen to reduce the likelihood of pre-existing CIN-tolerating mutations and to simplify the karyotype analyses. In cell lines that did not already have a p53 deletion, we homozygously deleted p53 so that they could continue to propagate despite ongoing chromosome missegregation (Extended Data Fig. 1c) (Santaguida et al. 2017). In addition, the vast majority of cancers with CIN have dysfunctional p53 pathways, making p53-deleted cells a good model for adaptation to CIN. The homozygous deletion of p53 in the cell lines led to no or very few karyotypic changes compared to wild-type counterparts, with the exception of HME1, which displayed the most initial chromosome aberrations (Extended Data Fig. 1d,e and Supplementary Table 2). Most cell lines showed a cell type-dependent decrease in their capacity to arrest at the G1/S transition following p53 deletion, as previously described (Extended Data Fig. 1f) (Kastan et al. 1991; Hartwell and Kastan 1994). Notably, the capacity of wild-type DLD1 and HME1 cell lines to arrest at G1 was generally low and did not significantly change upon p53 inactivation. In DLD1, we attribute this to the presence of the p53S241F mutation which fails to activate the p21 pathway (Sur et al. 2009). In HME1, this could be connected to the repression of the G1/S regulator p16INK4a, which has been described to occur in later passages of this cell type (Shapiro et al. 1998; Kim et al. 2002).

**Figure 1.**
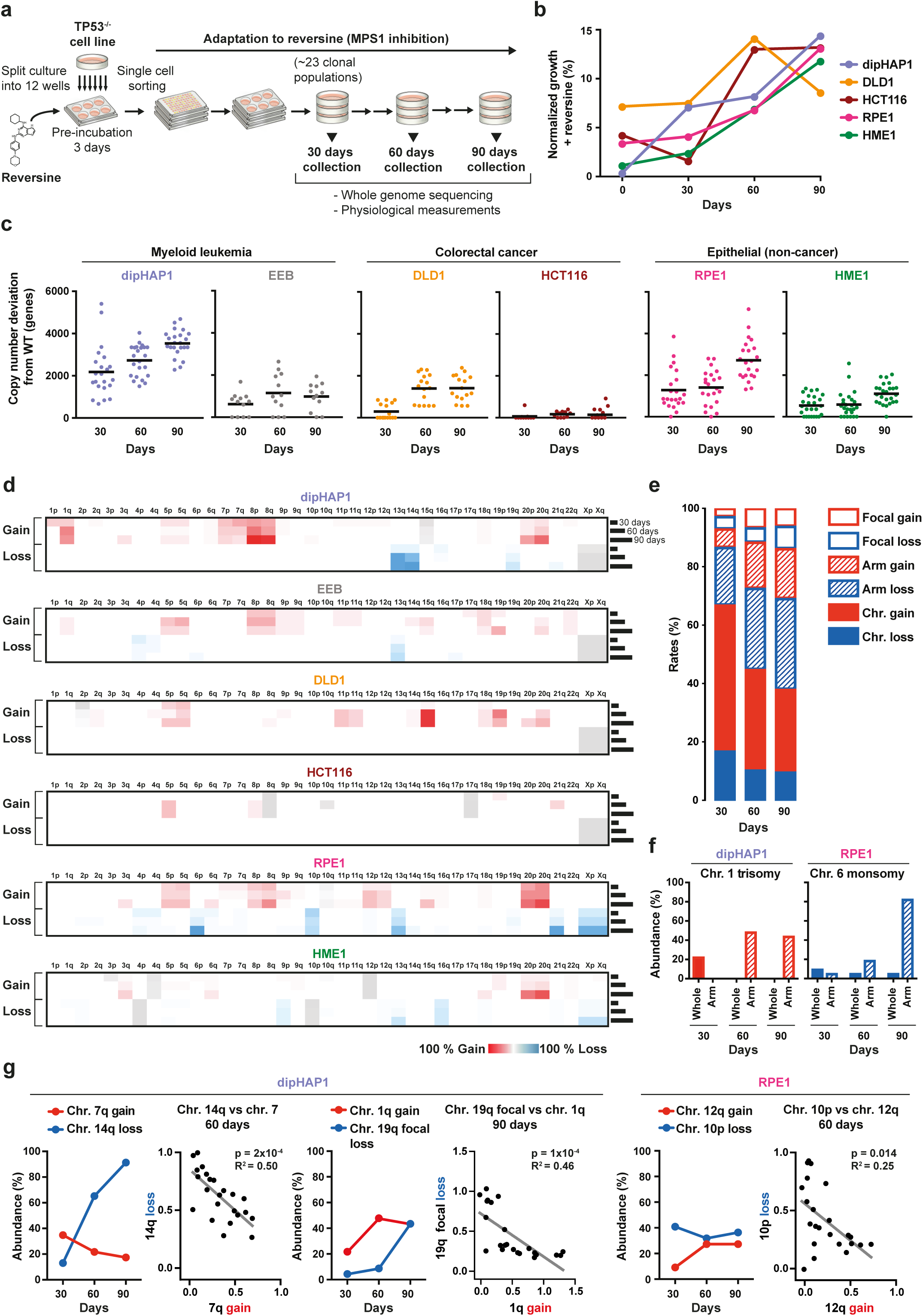
Long-term MPS1 inhibition leads to the selection of specific aneuploidy types and karyotype refinement over time. **a**, Schematic overview of the reversine adaptation process. **b**, Average of the growth in reversine of all populations of each cell line at 30, 60 and 90 days as measured using colony formation assays. Time point zero represents growth of unadapted parental cell lines. The percentages are normalized to the growth of the parental cell lines in the absence of reversine. **c**, Total copy number changes (of whole-chromosomes, arms and segments) of reversine adapted populations at 30, 60 and 90 days derived from Illumina sequencing read frequencies. Copy number alterations that were already present in the parental cell lines before adaptation are excluded. Black line represents the mean number of genes with altered copy numbers per adapted cell line. **d**, Heat-maps depicting the mean percentage of copy number changes in chromosome arms from all populations of each adapted cell line at 30, 60 and 90 days. Cell line inherent aneuploidies that were present at time point zero before the adaptation are colored in grey. Color code throughout the manuscript: red = chromosome gain and blue = chromosome loss. **e**, Quantification of the relative percentage of copy number changes of whole-chromosomes (filled), arms (dashed lines) and segments (focal changes, empty) across all reversine-cultivated cell lines at 30, 60 and 90 days. **f**, The percentage of adapted cell lines with whole and chromosome arm copy number changes of chr. 1 in dipHAP1 and chr. 6 in RPE1 respectively. **g**, Comparisons of the abundance of dipHAP1 or RPE1 adapted populations with specific chromosome losses or gains over time and the corresponding correlations between the chromosome copy number (0 = no loss/gain, 1 = chr. loss/gain; x-axis) for individual populations respectively at 60 or 90 days. p-values are from F-tests.

To obtain cycling cells adapted to long-term MPS1 inhibition, multiple cultures of the parental p53-deleted cell line were first pre-incubated in medium containing reversine for 3 days and then subjected to single cell sorting to obtain clonal populations. At least 20 independent clonal populations were then cultivated in the presence of reversine for a period of 90 days. In parallel, six populations from each cell line were propagated in the absence of reversine as a control. At each 30-day time interval, cell populations were analyzed using a combination of low-coverage whole-genome sequencing and fitness measurements. This simultaneous adaptation of many independent cultures allowed us to identify recurring karyotypic changes resulting from positive selection.

Most of the adapted cell populations showed a gradual increase in reversine resistance over time as measured by colony formation assays. Proliferation of cells in reversine increased from0.4-7.2 % relative to untreated cells before the adaptation (time point zero) to around 13-15 % at 90 days (Fig. 1b). Proliferation of the EEB cell line was not measured as the non-adherent nature of this cell line prevented colony formation. To determine the types of somatic copy number alterations acquired in the adapted cell lines, we measured copy number changes using low coverage next generation sequencing (NGS). Copy number was measured relative to the untreated parental population in 7.5 Mb intervals. From this analysis, we identified copy number changes of entire chromosomes, chromosome arms, and focal copy number alterations. We define aneuploidy as the gain or loss of either whole chromosomes or whole chromosome arms (Ben-David and Amon 2019). The degree of aneuploidy varied considerably between cell populations of the same cell type and even more so across cell types (Fig. 1c,d). On average, the dipHAP1 and RPE1 populations exhibited the highest degree of aneuploidy with 18 % and 14 % of the genome being aneuploid at 90 days, respectively. By contrast, the HCT116 populations showed the least amount of aneuploidy at ∼1 % of the genome on average. We conclude that aneuploidy formation in response to long-term MPS1 inhibition is highly cell line-dependent.

We next measured which types of copy number changes (whole chromosome, arm, or focal) were most prevalent in the adapted populations and how this changed over time. For all cell lines combined, whole-chromosome aneuploidies made up more than 60 % of all CNAs at the 30 days time point (Fig. 1e). Over time, the relative amount of whole chromosome aneuploidy decreased as arm and focal CNAs both increased. One reason for this trend can be seen in specific aneuploidies that changed from whole-chromosome CNAs at the 30 day time point into arm aneuploidies at later stages. This is exemplified by chr. 1 trisomy turning into the gain of only the q-arm in dipHAP1 cells and chr. 6 monosomy turning into monosomy of just the p-arm in RPE1 cells (Fig. 1f). The transition to smaller regions over time is also apparent from the increase in focal changes from 7 % at 30 days to 14 % at the 90 days time point. Notably, the increase in arm and focal aneuploidies was also seen when the acrocentric chromosomes were considered as whole chromosomes rather than chromosome arms (Extended Data Fig. 2a,b). The complete lack of whole chromosome and arm-level losses in DLD1 and HCT116, respectively, indicated fundamental differences in how the diverse cell lines adapted to the drug (Extended Data Fig. 2b). We conclude that early on in adaption to CIN, whole-chromosome changes are primarily acquired. At later stages, adapted populations develop more refined karyotypes by gaining and losing smaller chromosomal areas.

**Figure 2.**
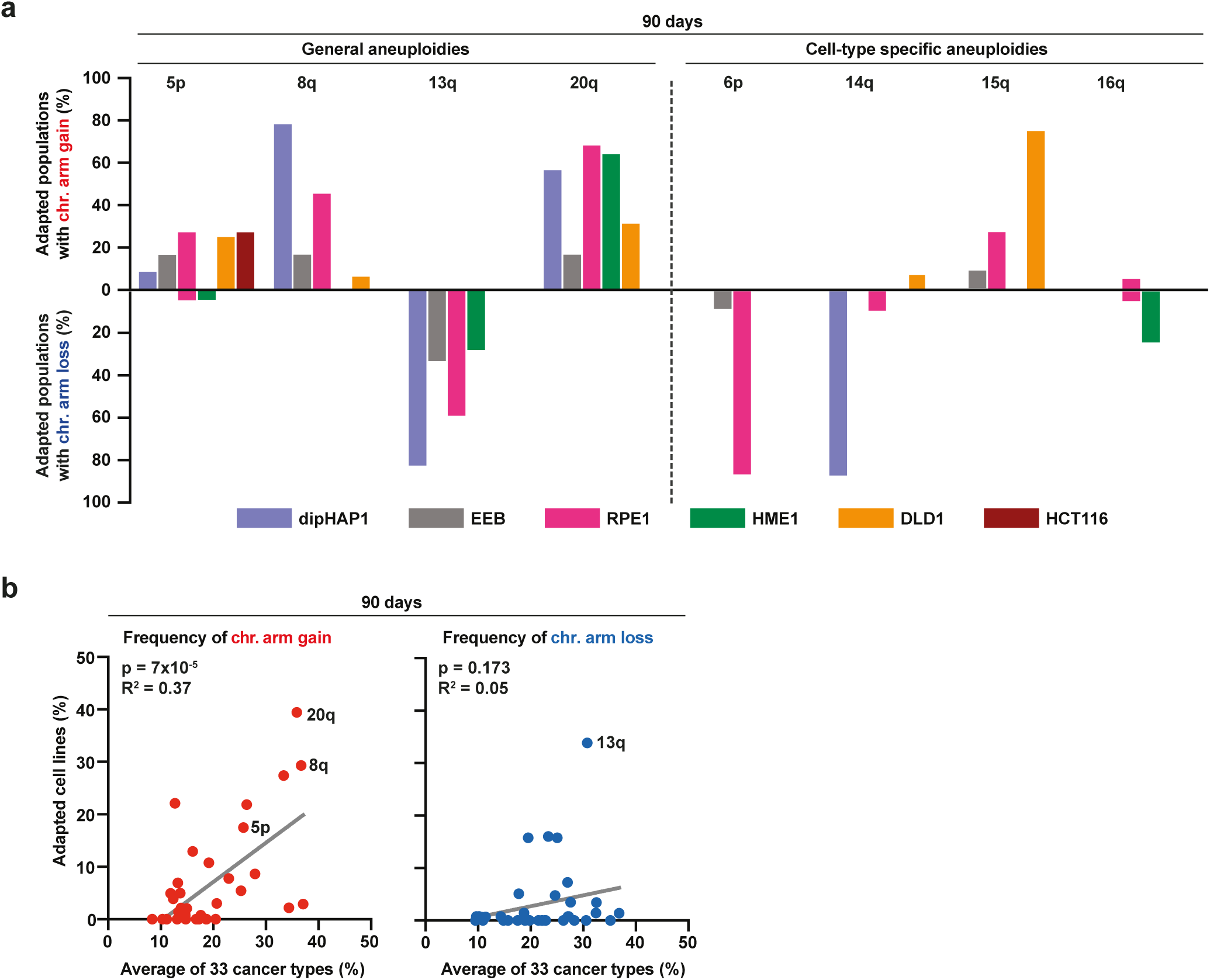
Reversine-cultivated cell populations converge on recurrent aneuploidy patterns that partially correlate with those seen in cancer. **a,** Percentage of adapted populations containing specific chromosome arm gains (upward) and chromosome arm losses (downward) among all populations of the different cell lines at the 90 days time point, **b,** Correlation between the frequencies of chromosome arm gains (left, red dots) and losses (right, blue dots) in reversine adapted cell populations after 90 days of adaptation and 33 different cancer types. The frequency of chr. arm gains or losses of the adapted cell lines represent the mean of the frequencies from all 6 different adapted cell lines, p-values are from F-tests.

### Correlations between aneuploidies over time

Although many aneuploidies increased in frequency over the course of the experiment, we were surprised to see that some aneuploidies that were frequently acquired early on in the adaptation process decreased in prevalence at later time points. These include the gain of chr. 7q and chr. 1q in dipHAP1 cells and the loss of chr. 10p in RPE1 cells (Fig. 1d,g). We hypothesized that these decreases could be due to incompatibilities with other, more beneficial chromosomes. For each of the examples mentioned above, we identified strong negative correlations with other aneuploidies that show corresponding increases over time. Aneuploidy of chr. 7q, 1q, and 10p, were each mutually exclusive with chr. 14q monosomy, chr. 19q partial loss, and chr. 12q gain, respectively (Fig. 1g). For chr. 19q, we observed a combination of whole arm monosomies and deletion of only the last 7.5 Mb of the chromosome (Extended Data Fig. 2c). The anticorrelation with chr. 1q gain was compared to both forms of 19q loss combined. These patterns indicate that distinct aneuploidies are lost when more beneficial aneuploidies take over the population. Indeed, for chr. 7q trisomy vs. chr. 14q monosomy and for chr. 10p monosomy vs. chr. 12q trisomy, the aneuploidy that emerged later in adaptation correlated better with growth in reversine (Extended Data Fig. 2d). We conclude that optimized aneuploid karyotypes develop over time through both the gradual selection of highly beneficial aneuploidies and the loss of less beneficial aneuploidies acquired early on in the adaptation process.

### Cell lines adapted to reversine converge upon recurrent aneuploidy patterns

We next determined which specific aneuploidies were most enriched at the end of the adaptation. For these analyses, we focused on whole chromosome and arm-level CNAs as these were most frequently observed. Cultivation of cell populations in the absence of reversine did not change karyotypes during the 90 days, with the exception of the loss of chr. Xp in dipHAP1 (Extended Data Fig. 3a). By time point 90 days, the vast majority of reversine-adapted populations had acquired recurring aneuploidies in all cell lines except for HCT116. Importantly, none of the cell populations underwent whole-genome duplication (WGD) during the 90 days of adaptation, demonstrating that WGD did not drive reversine resistance in these experiments (Extended Data Fig. 3b). There were some aneuploidies that were acquired in adapted populations across multiple cell lines. These include the gain of chromosomes 5, 8 and 20 in three or four out of the six adapted cell lines (Figs. 1d and 2a). In addition, chr. 13q was frequently lost in four cell lines. These results suggest that there are certain aneuploidies that provide a general benefit to reversine-induced CIN. In addition to the general aneuploidies, we also observed cell type-specific karyotypic changes. Examples of aneuploidies that only frequently occurred in one cell line include chromosome 15q gain in DLD1, 6p loss in RPE1, 14q loss in dipHAP1, and 16q loss in HME1. Each of these alterations was present in over 80 % of the independently adapted cultures at the 90 days time point (Fig. 2a). Notably, during the adaptation period, some populations reverted cell line-specific aneuploidies that were present at time point zero (Extended Data Fig. 3c). These include the loss of focal chr. 3p trisomy in HME1 and chr. X disomy in dipHAP1. The dipHAP1 cell line was generated through diploidization of male haploid chronic myeloid leukemia cells that lost chromosome Y. Therefore, dipHAP1 cells are aneuploid for chromosome X because they have two active copies. We conclude that both cell line-dependent and -independent aneuploidies are selected for during the adaptation to CIN.

**Figure 3.**
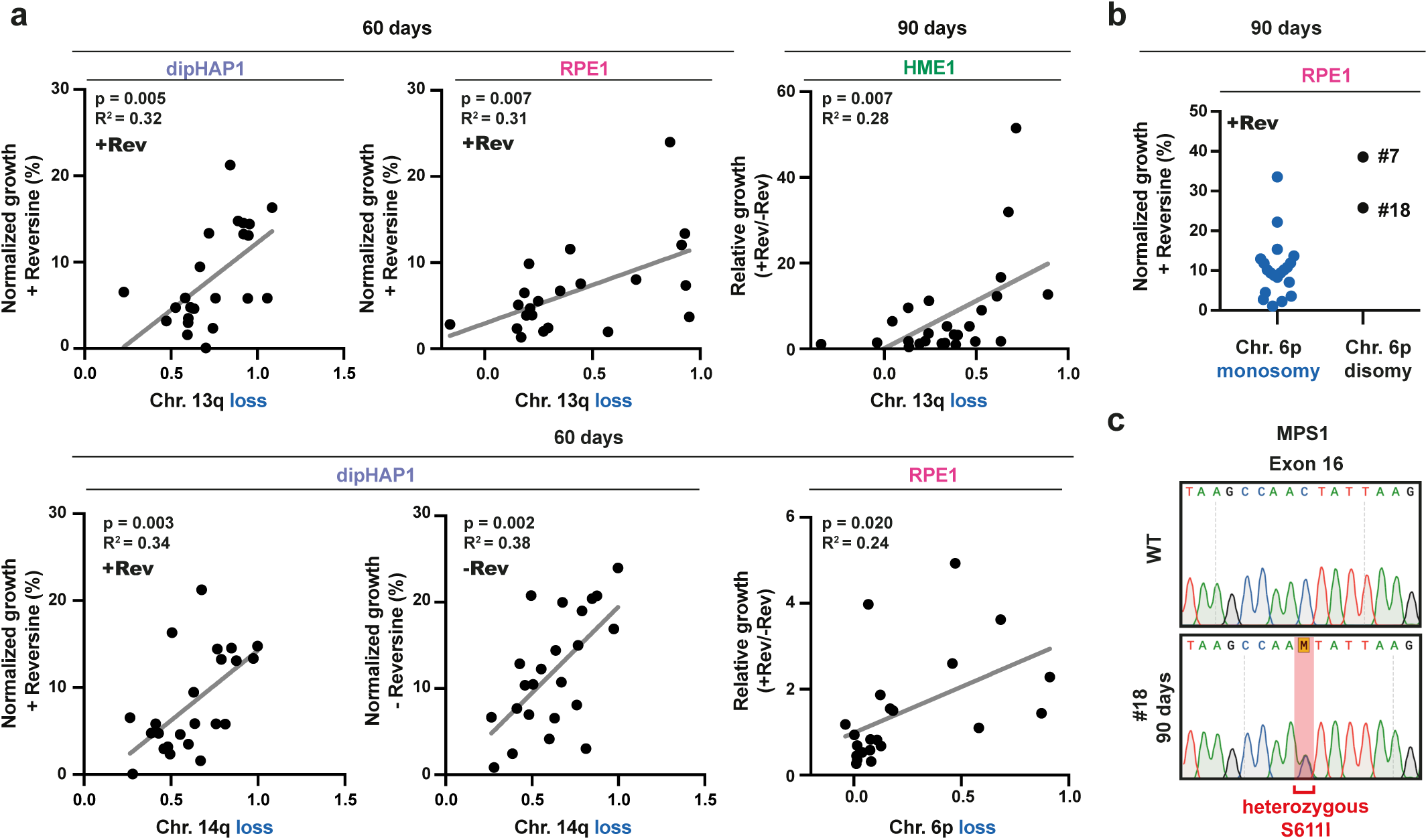
Specific arm aneuploidies correlate with better growth in reversine. **a,** Scatter plot of adapted populations from dipHAPI, RPE1 and HME1 cells harboring specific chromosome losses (0 = no loss, 1 = chr. loss; x-axis) vs. their normalized (+Rev or -Rev) or relative (+Rev/-Rev) growth (y-axis). Proliferation from colony formation assays was normalized to the untreated parental cell lines (WT). n = 2. Chromosome loss copy number data are from read frequencies using Illumina sequencing, p-values are from F-tests. **b,** Normalized growth in reversine of the 90 days adapted RPE1 populations grouped by chr. 6p copy number state. The two adapted RPE1 populations with 6p disomy are labeled with their cell line IDs #7 and #18. **c,** Sanger DNA sequencing chromatogram showing the presence of a heterozygous single base pair substitution in Exon 16 leading to the MPS1^S6111^ mutation in the RPE1 population #18 at the 90 days time point.

### The patterns of chromosome gains in reversine-adapted populations correlate with frequencies observed in cancer

To determine how similar aneuploid karyotypes of the adapted cultures were in comparison to cancer karyotypes, we analyzed copy number data from The Cancer Genome Atlas (TCGA) (The ICGC/TCGA Pan-Cancer Analysis of Whole Genomes Consortium 2020). Since we chose cell lines based on their lack of CIN and aneuploidy prior to the adaptation to reversine, the cancers that they are derived from (chronic myeloid leukemia and colorectal cancers with microsatellite instability) typically have low rates of aneuploidy (Hoadley et al. 2018). We therefore could not make direct comparisons between the aneuploid chromosomes present in the adapted cell lines and frequent aneuploidies in the corresponding cancer type. Instead, we combined the data of all of the reversine-adapted populations from the six cell lines and compared it to the combined data of 33 different cancer types from TCGA. At all three time points, chromosome gain frequencies between cancers and the adapted cell lines were highly correlated (Fig. 2b and Extended Data Fig. 3d). This pattern was largely driven by the high frequency of chromosome 8, 20, 5p, and 1q gains in both data sets. The only chromosome that is frequently gained across many different cancers but rarely in the adapted cell lines is chromosome 7. This difference may reflect the negative correlation between chr. 7q trisomy and other, even more beneficial aneuploidies under reversine selection as described above (Fig. 1g). These data suggest that our approach reconstituted fundamental chromosome gain patterns that are generally associated with proliferation advantages in cancer patients. It furthermore demonstrates that these advantages are largely independent of the underlying microenvironment, as they can be recapitulated by cell lines in culture. Curiously, the similarity between chromosome losses in the adapted cell lines and cancers was much weaker. This could indicate that the monosomies in the adapted populations are more specific to reversine resistance than general proliferation.

### Specific arm aneuploidies are associated with better growth in reversine

We next wanted to identify those aneuploidies that lead to reversine resistance. We therefore determined which of the recurrent arm aneuploidies in the adapted cell lines best correlated with improved growth in reversine. In addition to measuring growth in reversine, we also calculated growth in reversine relative to growth without the drug to account for potential negative effects on general fitness due to the aneuploidy burden. However, the complexity of the karyotypes at 90 days made the detection of significant correlations challenging. In addition, it was impossible to draw correlations in cases where aneuploidy of a particular chromosome was no longer observed. We therefore focused on looking for correlations with growth under reversine for aneuploidies that were found in 25-80 % of adapted populations at 90 days. For those with a frequency over 80 %, we looked at the 60 days time point when there were enough disomic populations for meaningful comparisons (Extended Data Fig. 4a,b).

**Figure 4.**
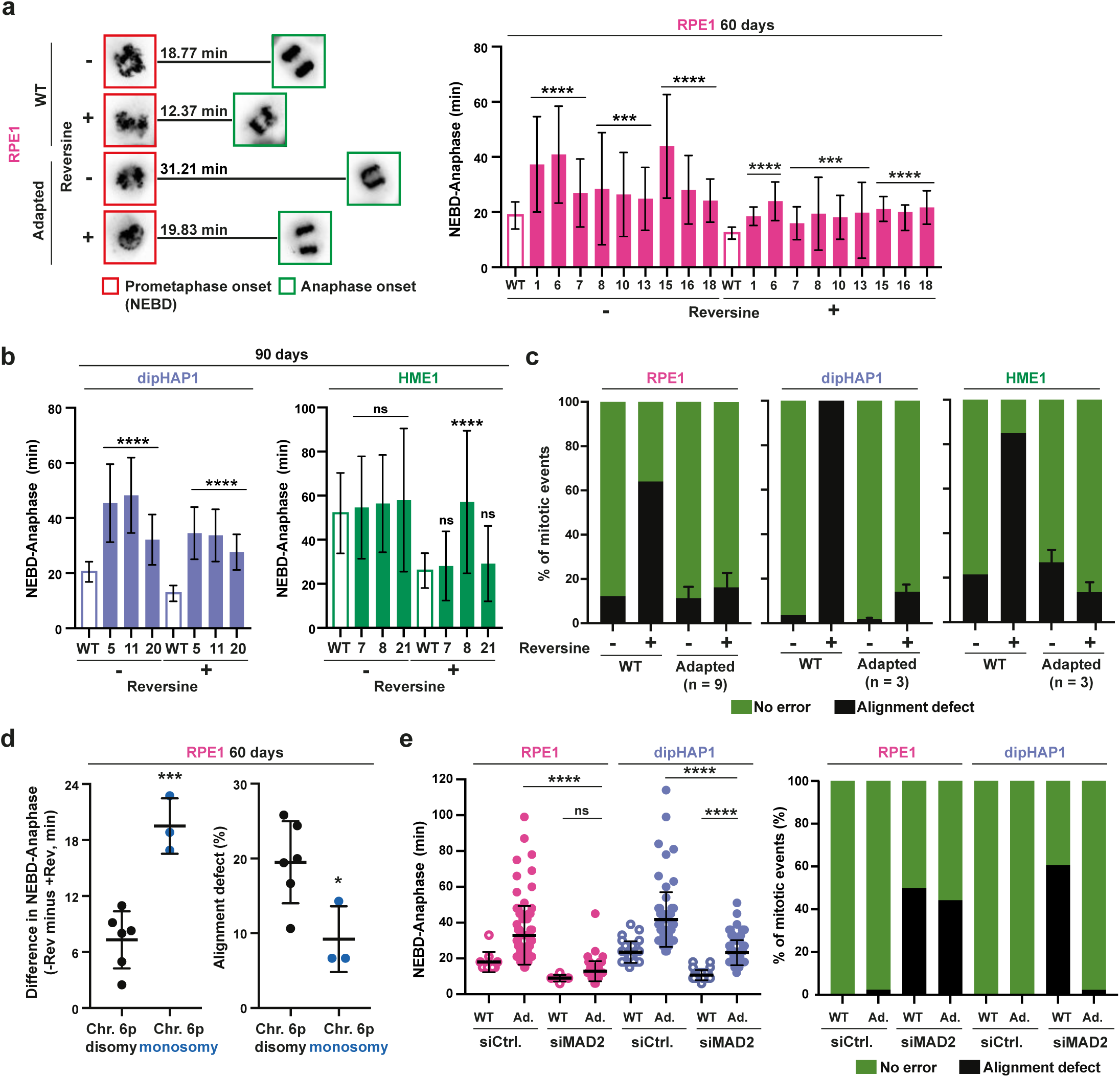
Adapted cells with specific aneuploidies limit CIN by increasing mitotic duration. **a**, Left: Representative images from a time-lapse of SiR-DNA-stained mitotic RPE1 cells at NEBD (red squares) and at anaphase onset (green squares) (full time-lapse is displayed in Extended Data Fig. 5b). The mean duration in minutes from NEBD to anaphase (n >25 mitoses) is shown. Right: Quantification of time from NEBD to anaphase (min) with and without reversine for the unadapted parental RPE1 cell line (WT, empty bar) and 9 adapted RPE1 cell lines (ID numbers, filled bars) at the 60 days time point. n = 3. **b**, Quantification as in **a**, only for dipHAP1 and HME1. Unadapted parental cell lines (WT, empty bar) and 3 adapted populations (ID numbers, filled bars) each are depicted. n = 3. **c**, Quantification of chromosome misalignment with and without reversine for the unadapted parental cell lines and the adapted cell populations from **a** and **b**. Alignment defects were measured as chromosomes that were not at the metaphase plate in the frame directly preceding anaphase. Error bars depict SD between the adapted populations (n depicted in graph) of the respective cell line. **d**, Left: Difference in the NEBD to anaphase timing for 60 days adapted RPE1 populations with and without reversine. Right: Percentage of mitoses with alignment defects with reversine in 60 days adapted RPE1 populations. Both classified by the copy number state of chromosome arm 6p. The cell lines are the same as in Figure **a**. **e**, Left: NEBD to anaphase duration for parental, 60 days adapted RPE1 and 90 days adapted dipHAP1 cell lines treated with either siRNAs for MAD2 or a scrambled control (siCtrl.). Ad. = adapted. Bars represent mean ± SD. p-values are from unpaired student’s t-tests, * = p<0.05, *** = p<0.001, **** = p<0.0001, ns = not significant.

We found positive correlations between reversine growth and 13q monosomy in both RPE1 and dipHAP1 cells at the 60 days time point and HME1 cells at the 90 days time point (Fig. 3a and Extended Data Fig. 4a,b). In addition, we observed positive correlations with chr. 6p loss in RPE1 cells and chr. 14q loss in dipHAP1 cells after 90 days of adaptation. However, the growth improvement of chr. 14q loss was just as strong in the absence of reversine, suggesting that the selection of this chromosome may not be reversine-specific. Positive correlations were also seen in both DLD1 and HME1 cells with 20q. However, in dipHAP1 cells, the gain of chr. 20 was negatively correlated with growth despite being present in ∼50 % of the adapted populations, emphasizing the difficulties of identifying the contributions of individual chromosomes in complex karyotypes. The two strongest candidates for aneuploidies that lead to a strong reversine resistance are therefore chr. 6p loss in RPE1 cells and chr. 13 loss in multiple cell lines.

### Point mutations that cause reversine resistance affect aneuploidy patterns

The correlation between reversine resistance and chr. 6p monosomy was significant at 60 days of adaptation, yet this trend was reversed at the 90 days time point (Fig. 3b). Surprisingly, after 90 days, the only two adapted RPE1 strains without chr. 6p monosomy had extremely high reversine resistance. We hypothesized that these cell lines adapted through point mutations instead, decreasing the need for beneficial aneuploidies. Since mutations in MPS1 had previously been shown to increase reversine resistance (Koch et al. 2016), we sequenced the kinase domain of ten of the adapted RPE1 cell lines. The only cell line with a mutation in MPS1 was indeed one of the two that did not obtain chr. 6p monosomy (Fig. 3c). The identified heterozygous S611I mutation is in the same amino acid that was previously demonstrated to confer resistance to reversine and multiple other MPS1 inhibitors (Koch et al. 2016). Since the MPS1 gene is not present on chr. 6p, we conclude that mutations present on one region of the genome can affect aneuploidy patterns in other regions of the genome.

### Adapted cells with specific aneuploidies rescue reversine treatment by increasing the duration of mitosis

Next, we investigated if increased reversine resistance was associated with decreased mitotic errors in the adapted cell lines. For this, we monitored mitotic timing and chromosome segregation fidelity by live cell imaging of unsynchronized populations stained with SiR-DNA (Fig. 4a,b and Extended Data Fig. 5a,b). The analysis comprised 15 adapted cell populations (9 from RPE1 time point 60 days and 3 from dipHAP1 and 3 HME1 time point 90 days) that showed extensive karyotype diversity in the most commonly acquired aneuploidies: 6p loss, 13q loss and 20q gain.

**Figure 5.**
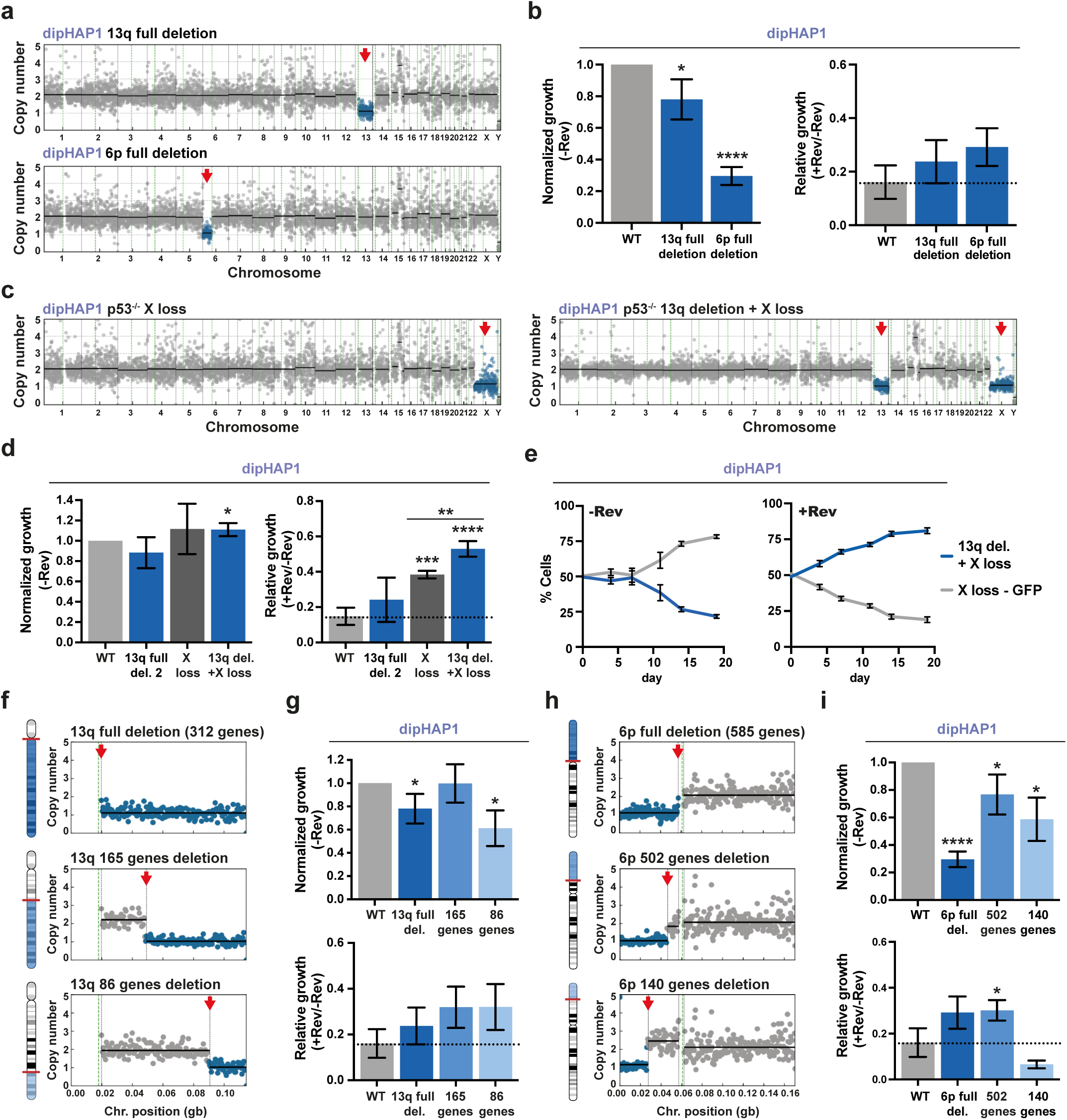
Engineered individual monosomies and partial chromosome deletions can recapitulate reversine resistance. **a**, Copy number profiles of engineered dipHAP1 cell lines with the deletion of chromosome arm 13q or 6p. Blue dots represent region of chromosome loss which is highlighted with a red arrow. **b**, Normalized (-Rev) and relative (+Rev/-Rev) growth of the depicted 13q and 6p deletion cell lines. Proliferation measurements are from colony formation assays. Normalized growth (-Rev) represents the growth of the cell line without reversine normalized to the proliferation of the dipHAP1 parental cell line (WT) without reversine. The relative growth (+Rev/-Rev) represents the proliferation of the cell line with reversine (+Rev) relative to its growth without reversine (-Rev). The dotted line depicts mean relative growth of dipHAP1 WT cells with reversine. n = 3. **c**, Copy number profiles of engineered dipHAP1 cell lines with the loss of chromosome X and chr. 13q deletion in addition to X. **d**, Normalized and relative growth of the depicted 13q, X, or double monosomic cell line. For WT and 13q del. 2: n = 6; for X loss and 13q del. + X loss: n = 3. **e**, Co-cultivation of a GFP-labeled cell line with the monosomy of chr. X and a cell line with monosomies of both X and 13q with and without reversine. The mean +/-SD for 6 separate populations of cells in one time course is shown. **f**, Representative depiction and chromosome-specific copy number profiles of engineered partial deletions of chromosome arm 13q. **g**, Normalized and relative growth of the 13q partial deletion cell lines. n = 3. **h**, Representative depiction and chromosome-specific copy number profiles of engineered partial deletions of chromosome arm 6p. **i**, Normalized and relative growth of the 6p partial deletion cell lines. n = 3. Bars depict mean ± SD. p-values are from unpaired student’s t-tests, * = p<0.05, ** = p<0.01, *** = p<0.001, **** = p<0.0001. Where not indicated, the p-value was greater than 0.05.

Elongation of mitotic duration has been previously proposed as a resistance mechanism to MPS1 inhibition (Sansregret et al. 2017). Although the three parental cell lines had substantial differences in mitotic duration (∼19 min in RPE1 and dipHAP1 cells vs. ∼50 min in HME1 cells), the average duration from nuclear envelope breakdown (NEBD) to anaphase was reduced after reversine addition by ∼6 to 20 minutes (Fig. 4a,b) (Santaguida et al. 2010). Strikingly, for the adapted cell lines, the mitotic timing in the presence of reversine was now generally more similar to the timing of unadapted cells (WT) in the absence of reversine. This suggests that cells may adapt to reversine by extending the mitotic duration to provide additional time for chromosome alignment. Notably, upon reversine removal, mitotic timing became exceedingly long in adapted cells with a fraction of mitoses being longer that 120 min (Extended Data Fig. 5c).

The shortened mitotic timing after reversine treatment greatly increases the number of misaligned chromosomes. To determine if the adapted cells had improved chromosome alignment, we measured the percentage of cells with one or more chromosomes that where not properly aligned at the metaphase plate immediately before anaphase onset. Adapted cells exhibited very low chromosome mis-alignment rates in the presence or absence of reversine (Fig. 4c). For RPE1 cells, we identified correlations between the duration of mitotic timing with reversine and both the relative growth in reversine and the number of alignment errors (Extended Data Fig. 5d,f). In addition to misaligned chromosomes, reversine addition also substantially increases the formation of lagging chromosomes and chromosome bridges in RPE1 and dipHAP1 cells. The adapted cell lines display a decrease in the rates of anaphase errors in the presence of reversine (Extended Data Fig. 5e). These results indicate that the adapted cells limit reversine-induced mitotic error rates by extending mitotic duration.

After having determined that the adapted cell lines display an increased mitotic duration and decreased CIN in reversine, we next asked if these traits were associated with any recurrent aneuploidies. We found that loss of chr. 6p, which is associated with better relative growth in reversine (Fig. 3a and Extended Data Fig. 4b), is also associated with increased mitotic length and decreased chromosome mis-alignment rates (Fig. 4d).

Since MPS1 activates the SAC and we used reversine concentrations that do not completely inhibit its activity, cells could adapt either by increasing the residual SAC activity or by inhibiting activities downstream of the SAC such as the APC/C. We hypothesized that in reversine-adapted cells, SAC and/or APC/C activity could be altered to partially restore mitotic timing in the presence of the drug. To test this, we used small interfering RNA (siRNA) to deplete the essential SAC component MAD2 in RPE1 and dipHAP1 parental and adapted cell lines (Extended Data Fig. 6a). Similar to reversine treatment, MAD2 depletion in unadapted cells led to a decrease in mitotic length and increased alignment errors (Fig. 4e and Extended Data Figure 6b). MAD2 depletion eliminated the mitotic delay phenotype exhibited by adapted RPE1 cells in the absence of reversine. Upon SAC inactivation, both unadapted and adapted RPE1 cells became extremely similar in mitotic duration and chromosome mis-alignment rates. These results indicate that in adapted RPE1 cells, the activity of the SAC is modulated to counteract the effects of partial MPS1 inhibition. In contrast to RPE1 cells, mitotic length in dipHAP1 cells was still significantly higher than in unadapted cells after MAD2 depletion. In addition, MAD2 depletion in the adapted dipHAP1 cells did not increase chromosome mis-alignment rates. These partial phenotypes could be either due to dipHAP1 cells adapting through mechanisms downstream of the SAC, or only partial inhibition of SAC activity by the MAD2 depletion. However, nocodazole treatment led to a fully penetrant mitotic arrest in control cells and caused mitotic slippage in 100 % of MAD2-depleted RPE1 and dipHAP1 cells, indicating robust SAC inactivation (Extended Data Fig. 6b). We conclude that RPE1 cells adapt to reversine primarily by promoting SAC activity, while dipHAP1 cells adapt through a combination of increased SAC activity and additional effects downstream of the SAC such as the APC/C.

**Figure 6.**
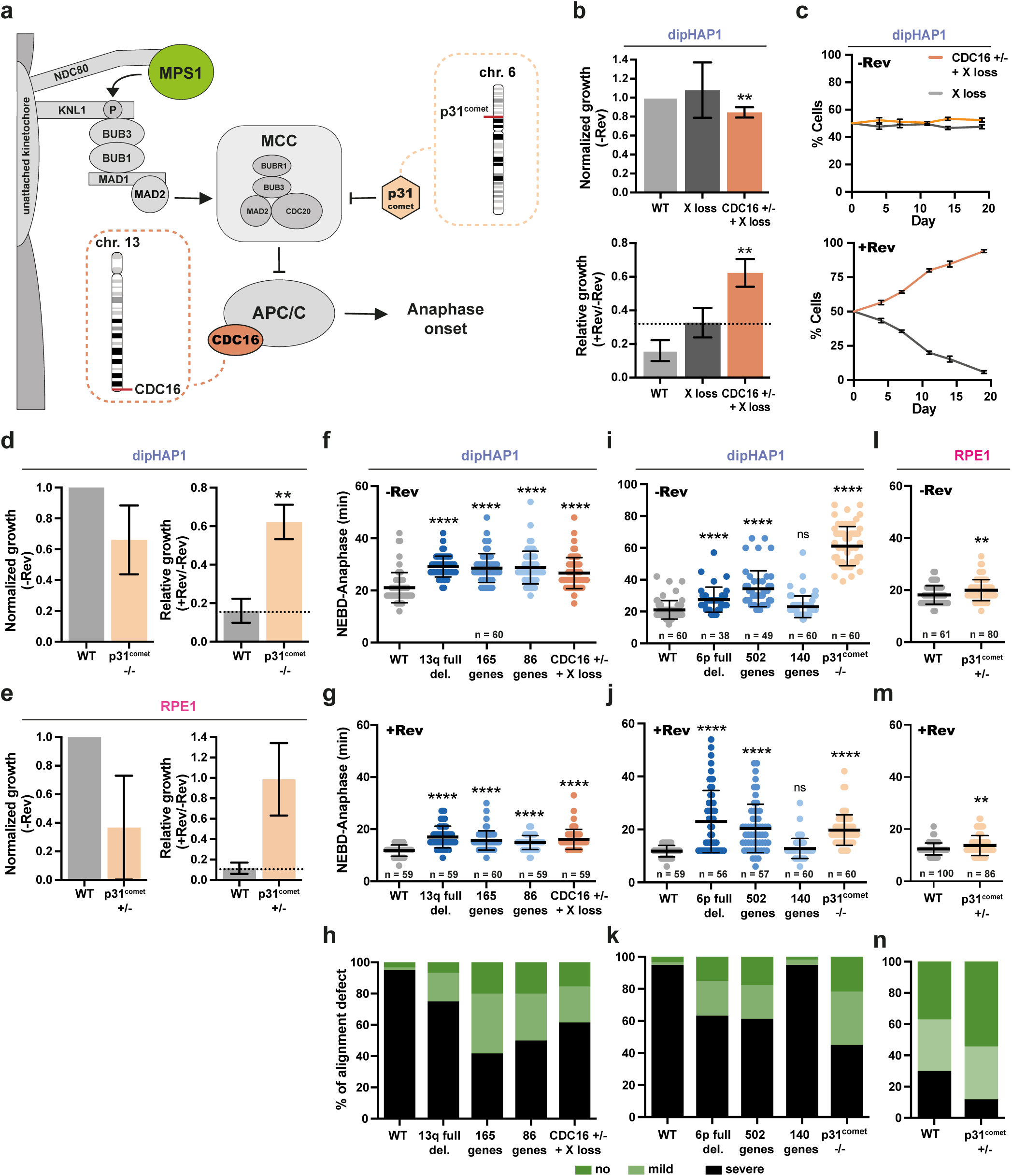
Identification of the cellular mechanism underlying reversine resistance. **a**, Simplified model of the SAC and the roles of CDC16 and p31^comet^. **b**, Normalized and relative growth of the dipHAP1 CDC16 heterozygous knockout (+/-, + X loss) and a X loss cell line as control. Proliferation measurements are from colony formation assays. Normalized growth (-Rev) represents the growth of the cell line without reversine normalized to the proliferation of the dipHAP1 parental cell line (WT) without reversine. The relative growth (+Rev/-Rev) represents the proliferation of the cell line with reversine (+Rev) relative to its growth without reversine (-Rev). The dotted line represents mean relative growth of the X loss cell line. n = 3. **c**, Co-cultivation of the GFP-labeled X loss cell line and a cell line with CDC16 heterozygous knockout and monosomy of chromosome X with and without reversine. The mean +/-SD for 6 separate populations of cells in one time course is shown. **d,** Normalized and relative growth of the parental dipHAP1 and p31^comet^ homozygous knockout cell lines. n = 2. **e**, Normalized and relative growth of the parental RPE1 and p31^comet^ heterozygous knockout cell lines. n = 2. **f,** Duration from NEBD to anaphase in min for the engineered dipHAP1 13q deletions and the CDC16 +/-knockout cell line without reversine (-Rev), (n = 60 mitoses) and **g**, with reversine (+Rev), (n = 59-60 mitoses). **h,** Quantification of no (dark green), mild (one unaligned chromosome, light green) or severe (multiple unaligned chromosomes, black) alignment defects of the mitoses from the dipHAP1 parental, 13q partial deletion, and the CDC16 +/-knockout cell lines. **i**, Duration from NEBD to anaphase in min for the engineered dipHAP1 6p deletions and the p31^comet^ -/-knockout cell line without (-Rev), (n = 38-60 mitoses) and **j**, with reversine (+Rev), (n = 56-60 mitoses). SiR-DNA staining efficiency was increased by adding 10 µM Verapamil. **k**, Quantification of the alignment defect of the measured mitoses from the parental, 6p partial deletion, and the p31^comet^ -/-knockout cell lines as in **h**. **l**, Duration from NEBD to anaphase in min for the RPE1 p31^comet^ +/-knockout cell line without reversine (-Rev), (n = 61-80 mitoses) and **m**, with reversine (+Rev), (n = 86-100 mitoses). **n**, Quantification of the alignment defect of the parental RPE and p31^comet^ +/-knockout cell lines as in **h,k**. For graphs f-k: n = 3; for l-n: n = 5. Bars depict mean ± SD. p-values are from unpaired student’s t-tests, ** = p<0.01, *** = p<0.001, **** = p<0.0001, ns = not significant.

### Engineered individual monosomies are sufficient for partial reversine resistance

Next, we sought to identify the basis behind the frequent selection of specific aneuploidies in the adapted cell lines. We focused on the most common chromosome losses, as we were able to efficiently engineer specific chromosome and gene losses as opposed to gains. We chose chromosome arm 13q loss, since it is one of the most frequently selected aneuploidies across all reversine adapted cell lines (Fig. 2a). In addition, we decided to examine monosomy of chr. 6p since it correlated well with improved reversine resistance in RPE1 cells (Fig. 3a and Extended Data Fig. 4b). To ascertain the specific roles of these monosomies in reversine resistance we engineered the losses of chr. arms 6p and 13q individually in the dipHAP1 cell line with CRISPR (Extended Data Fig. 7a).

To generate cell lines with one specific chromosome arm loss, we selected sgRNAs that target the chromosome arm either at one site close to the centromere or at a repetitive region in the chromosome arm (Zuo et al. 2017) (Extended Data Fig. 7b and Supplementary Table 3). Two different sgRNAs for each chromosome arm deletion were used to account for off target effects of the individual sgRNAs. Using this approach, we successfully engineered two 13q monosomy and three 6p monosomy cell lines in dipHAP1 cells. These cell lines contain no other CNAs (Fig. 5a and Extended Data Fig. 7c). Attempts to engineer chromosome losses in RPE1 cells were unsuccessful, potentially due to differences in DNA repair mechanisms.

In the absence of reversine, engineered dipHAP1 cells monosomic for chromosome arm 13q showed a mild growth reduction, whereas deletion of one copy of 6p resulted in very poor growth (Fig. 5b and Extended Data Fig. 7d,e). Both monosomies showed a better growth in reversine relative to its absence when compared to wildtype dipHAP1 cells as measured by colony formation assays. However, for the 6p deletion, the added reversine resistance could not overcome the negative effect of the aneuploidy on proliferation, which may explain why this CNA was not observed in the adapted dipHAP1 cells.

Occasionally, other aneuploidies were observed during the screening process for 6p and 13q loss. These include cell lines whose only CNA is the loss of chromosome X, Xp, and 19p (Extended Data Fig. 7f). As mentioned earlier, the loss of chr. X results in a relief from aneuploidy in dipHAP1 cells and was observed in all of the reversine-adapted populations (Extended Data Fig. 3c). Similar to 6p and 13q monosomy, the loss of the whole X chromosome showed an increased reversine resistance compared to euploid wild type cells (Extended Data Fig. 7g). Loss of only the p-arm of X did not show this phenotype, suggesting that the gene(s) affecting reversine resistance are on the q-arm. Loss of 19p was not observed in any of the adapted cell lines. We therefore used this aneuploidy as a control for the general effects of chromosome loss. Chromosome 19p loss made the cells more sensitive to reversine, demonstrating that reversine resistance is not a general property of monosomies (Extended Data Fig. 7h). We conclude that the loss of one copy of chromosomes 6p, 13q, or X is sufficient to confer reversine resistance.

Notably, the rescue phenotypes for the engineered monosomies were much weaker than what was observed in the adapted cell lines, suggesting that multiple aneuploidies act cooperatively during the adaptation process. To test if multiple monosomies can act together to create stronger drug resistance phenotypes, we used a chr. 13q deletion cell line that also acquired chr. X loss during the generation of the cell line (Fig. 5c). The combined double monosomic cell line had substantially greater reversine resistance than either monosomy individually as measured using colony formation assays (Fig. 5d). To additionally measure reversine resistance with a more sensitive assay, we performed a co-cultivation assay between a GFP-labeled cell line with chr. X monosomy and an unlabeled cell line with chr. X and chr. 13q monosomy (Extended Data Fig. 7i,j) In the absence of reversine, the cell line with only a single aneuploidy grew better, but with the addition of reversine, the double monosomic cell line quickly and reproducibly took over the population (Fig. 5e and Extended Data Fig. 7k). These results demonstrate that multiple monosomic chromosomes can create stronger phenotypes and supports the idea that the reversine resistance in the adapted cell lines results from the additive or synergistic contributions of multiple aneuploid chromosomes.

### Identification of chromosomal regions that are sufficient to confer reversine resistance

The losses of chromosome arms 6p and 13q decrease the copy number of hundreds of genes on the aneuploid chromosome. To narrow down the chromosomal regions that confer drug resistance, we engineered partial deletions of the chromosome arms. We engineered two partial deletions that eliminate either 165 or 86 of the 312 genes on chr. 13q (Fig. 5f and Extended Data Fig. 8a). Both of these deletions exhibited robust reversine resistance that was similar to the full arm deletion (Fig. 5g and Extended Data Fig. 8b). This indicates that either one or more of the 86 genes closest to the telomere contribute to reversine resistance when heterozygously deleted or another genetic element in this region is responsible. In addition to full arm loss, we also detected focal losses on chr. arm 13q in the adapted dipHAP1 and HCT116 cell lines (Extended Data Fig. 8c). Focal losses of chr. arm 13q in the adapted cell lines always contained our identified chromosomal region, further supporting the results from the engineered partial deletions.

The full deletion of chromosome arm 6p decreases the copy number of 585 genes. We again made two partial deletions, one that eliminates 502 genes and another that only removes 140 genes (Fig. 5h and Extended Data Fig. 8d). Interestingly, although both of the smaller deletions grew better in the absence of reversine than the full arm deletion, only the larger 502 gene deletion still showed better growth in reversine than without reversine relative to the parental cell line (Fig. 5i and Extended Data Fig. 8e). This narrows down the region of interest to the one in between the partial deletion sites containing 362 genes. Importantly, these results further demonstrate that the reversine resistance is due to the loss of specific regions and not a general effect of aneuploidy or the technique used for aneuploidy generation.

### Identification of genes that confer reversine resistance when heterozygously deleted

Monosomy of chromosomes can induce phenotypes either through loss of heterozygosity (LOH) or decreased expression of genes on the chromosome. Since the monosomies were engineered in the dipHAP1 cell line that is already homozygous for nearly the entire genome (with the exception of a portion of chromosome 15), we can eliminate LOH as a possible contributor to the aneuploidy phenotypes. After further refining the regions of interest on chr. 13q and 6p, we took a candidate approach to identify genes that may contribute to the phenotype. The 86 gene region on 13q contains the gene for CDC16, an essential component of the anaphase promoting complex/cyclosome (APC/C) (Fig. 6a). A mutation in CDC16 was previously identified as conferring reversine resistance, making it a strong candidate for a gene that could contribute to resistance when heterozygously deleted (Sansregret et al. 2017). We therefore used CRISPR to engineer a heterozygous deletion of CDC16 in dipHAP1 cells (Extended Data Fig. 9a). Whole genome sequencing of the identified cell line revealed that it also contained a deletion of chromosome X (Extended Data Fig. 9b). We therefore used the chr. X deletion cell line described above as the control for experiments with this cell line. CDC16 +/-cells showed a strong resistance to reversine, substantially greater than the resistance observed for X loss alone (Fig. 6b). CDC16 Western blot analysis of the deletion and knockout cell lines further revealed that heterozygous deletions can impact protein levels (Extended Data Fig. 9c). Additionally, co-cultivating the CDC16 knockout cell line with a GFP-labeled X-loss control cell line showed that the knockout cell line quickly overtook the population in medium containing reversine in two independent experiments (Fig. 6c and Extended Data Fig. 10a). This level of resistance is consistent with the CDC16 gene being primarily or solely responsible for the reversine resistance in the chr. 13q deletion cell lines, indicating that the loss of CDC16 is likely the driving force behind the selection of chr. 13q monosomy in the adapted cell lines.

The 362 gene region on chr. 6p contains two genes of primary interest for potentially contributing to reversine resistance, p31^comet^ and p21^CDKN1A^. p31^comet^ is a negative regulator of the mitotic checkpoint complex (MCC) that was previously identified in an CRISPR/Cas9 screen for MPS1 resistance (Fig. 6a) (Thu et al. 2018). p21^CDKN1A^ is a tumor suppressor gene that promotes cell cycle arrest following chromosome missegregation (Karimian, Ahmadi, and Yousefi 2016). Although p21^CDKN1A^ frequently acts through p53, it also has p53-independent functions that might contribute to reversine resistance in our p53-deleted cell lines (Matsuda et al. 2017). However, neither heterozygous nor homozygous deletion of p21^CDKN1A^ conferred reversine resistance in dipHAP1 cells (Extended Data Fig. 9d,e,f). For p31^comet^, we obtained a homozygous deletion cell line in dipHAP1 cells and a heterozygous deletion cell line in RPE1 cells (Extended Data Fig. 9g,h). Both cell lines showed strong resistance to reversine (Fig. 6d,e). p31^comet^ protein abundance was proportional to the gene copy in the engineered cell lines (Extended Data Fig. 9i). We conclude that the heterozygous deletion of p31^comet^ is sufficient for robust reversine resistance, and that the decrease in p31^comet^ expression is likely the principal basis behind the selection of 6p loss in RPE1 cells adapted to reversine.

### Characterization of the cellular mechanisms behind the reversine resistance

The adapted dipHAP1 and RPE1 cell lines showed an increased time in mitosis compared to the parental cell lines (Fig. 4a,b). In addition, they displayed improved chromosome alignment in the presence of reversine (Fig. 4c). We next determined if the engineered monosomies and heterozygous knockout cell lines that induce reversine resistance also rescue the cell cycle timing and chromosome alignment defects.

The chr. 13q monosomy, the partial chr. 13 deletions, and the CDC16 heterozygous deletion all had nearly identical NEBD to anaphase delays of ∼8 minutes in the absence of reversine (Fig. 6f). When reversine was added, the mitotic timing was reduced for all of the mutants, yet still significantly higher than for the parental cell line (Fig. 6g). Similarly, all of the cell lines affecting chr. 13 partially rescued the reversine-induced chromosome alignment defect (Fig. 6h). The chr. 6p full and the 502 gene deletions also had a highly significant increase in mitotic duration both with and without reversine and a rescue of chromosome alignment (Fig. 6i-k). In addition, the p31^comet^ homozygous deletion in dipHAP1 cells resulted in extremely long mitotic delays with and without reversine and a robust rescue of the alignment defects. By contrast, the 140 gene partial deletion that did not rescue reversine resistance did not show an anaphase delay nor a rescue of chromosome alignment. In RPE1 cells, the p31^comet^ +/-cell line had a small but significant anaphase delay in both the presence and absence of reversine (Fig. 6l,m). An improvement in chromosome alignment was also observed in this cell line (Fig. 6n). As with the rescue of proliferation in reversine, these phenotypes were not as strong as what was observed for the adapted cell lines, further suggesting that the adaptation is due to the cumulative effects of multiple genomic changes.

A recent study suggests that the presence of monosomic chromosomes generally increases mitotic duration in RPE1 cells (Chunduri et al. 2021). To determine if the elongation of mitosis is a general trait of our engineered monosomies, we once again used the dipHAP1 cell line with the loss of chr. 19p (591 genes). This cell line showed no differences in mitotic timing with or without reversine and no rescue of the chromosome alignment defects (Extended Data Fig. 10b,c). We conclude that the mitotic delay is not caused by monosomy in general in dipHAP1 cells, and that the observed phenotypes are chromosome dependent. Overall, these results are consistent with the reversine growth data, demonstrating a clear link between mitotic timing, chromosome alignment, and drug resistance for the engineered aneuploid and gene knockout cell lines.

## Discussion

In this study, we developed a time-resolved adaptation assay for human cells to study complex karyotype formation over time in response to long-term MPS1 inhibition in six different cell lines. Reversine-mediated inactivation of the SAC leads to high rates of CIN that also serve as a selective force to adapt via aneuploidy (Fig. 1a). Thus, our strategy is comparable to one that we previously used to generate complex aneuploidy in yeast (Ravichandran et al. 2018). Similar to what was observed in yeast, independently cultivated populations became more homogenous over time and converged on optimal karyotypes.

During the development of these optimized karyotypes, we observed different types of patterns that bear a striking similarity to those previously reported in cancer cells. Some of the most frequently obtained aneuploidies were observed across many of the adapted cell lines. This is analogous to certain aneuploidies being frequently observed in the majority of cancer types such as the gain of chromosomes 8 or 20. By contrast, other aneuploidies in the adapted cell lines were only observed in a single cell line. These included 6p loss in RPE1 and 16q loss in HME1. Such cell line-specific aneuploidy patterns reflect cancer type-specific aneuploidies, which include chromosome 16 loss in breast and ovarian cancers and chromosome 3p gain specifically in sarcomas (Hoadley et al. 2018). This cell-line specificity could be either due to differential gene expression, interactions with other common mutations, or both. In support of the second hypothesis, we found a mutation in the MPS1 gene specifically in a cell line with a karyotype that was highly divergent from the other adapted populations. Altered aneuploid karyotypes resulting from specific mutations has also recently been reported in yeast (Clarke et al. 2022). In conclusion, the ability to recapitulate both general and cell line-specific aneuploidy patterns in human cells is an important step in determining the bases behind these patterns. Moreover, we found a strong correlation between chromosome gain patterns observed in our CIN-adapted cell populations and those frequently observed in cancer, including the gains of chromosomes 1q, 5p, 8, and 20. This suggests that the conditions for acquiring chromosome gains are independent of microenvironment and instead reflect more general cell autonomous proliferation advantages through alterations in the cell cycle machinery. This result is in agreement with the proposed roles for these aneuploidies in promoting proliferation through the upregulation of genes involved in cell cycle entry and DNA repair (Su et al. 2021; Dehner et al. 2021; Tabach et al. 2011; Scotto et al. 2008).

In addition, we observed that smaller CNAs become more prevalent over time, with an increase in arm-level and segmental copy number changes relative to whole chromosome aneuploidies. Since the benefits of aneuploidy come at the cost of expression imbalances of the other genes present on the aneuploid chromosome, reducing the number of imbalanced genes would be advantageous. The increase in arm aneuploidy frequencies over time has also been observed in cancer karyotype analyses (Shukla et al. 2020). Our observations suggest a basis for this trend, in which whole chromosome aneuploidies are converted to arm-level or smaller CNAs over time within a population to partially relieve the aneuploidy burden.

Further, we observe strong negative correlations between specific aneuploid chromosomes. Such correlations are also observed in cancer karyotypes (Ravichandran et al. 2018; Shukla et al. 2020). Over time, the more beneficial aneuploidy replaces the less beneficial one, such as with the transition from chr. 7 trisomy to chr. 14 monosomy in dipHAP1 cells. Importantly, this type of negative pattern would not be observable by simply looking at the karyotypes at later time points, such as in fully developed cancers. The replacement of one aneuploid chromosome with another may also provide the basis behind the frequent chromosome-level copy-number neutral loss of heterozygosity in cancer (Nichols et al. 2020; Ciani et al. 2022). In yeast, these negative correlations can result from genetic interactions between the aneuploid chromosomes, which may also be the basis behind them in human cells (Ravichandran et al. 2018).

The final similarity between the aneuploidy patterns we observe and cancer karyotypes is that the HCT116 and DLD1 colorectal cell lines used in this study that exhibit microsatellite instability (MSI) acquire very few chromosome gains and no chromosome losses. This observation is in line with the mutual exclusivity between MSI and CIN in colorectal cancers (Lengauer, Kinzler, and Vogelstein 1997; Eshleman et al. 1998). The complete lack of chromosome losses in these cell lines may reflect the accumulation of mutations that recapitulate loss-of-function phenotypes similar to chromosome losses. By contrast, the increased expression caused by chromosome gains may be more difficult to obtain via mutations acquired from a loss of mismatch repair, which may explain why the DLD1 cell line still acquires chromosome gains. Overall, the ability to recapitulate these types of aneuploidy patterns in human cells through adaptation experiments gives insights into the bases behind their formation and provides a handle for directly testing the driving forces behind many of the most prominent motifs observed in cancer karyotypes.

Correlations between growth in reversine and specific CNAs in the adapted cell lines implicated 13q loss in multiple cell lines and 6p loss in RPE1 cells with resistance to the drug. In addition, chr. 6p monosomy was correlated with increased mitotic timing and improved chromosome alignment in the presence of reversine. Engineered monosomies of chromosome arms 13q and 6p recapitulated reversine resistance, extended mitotic timing, and decreased mitotic errors. Downregulation of p31^comet^ or CDC16 through the monosomies of chromosome arms 6p or 13q should both lead to decreased APC/C activity, which would delay anaphase onset. It has previously been shown that the deletion of the APC/C subunits APC7 or APC16, which also should lead to a decreased APC/C activity, rescues the lethality of MAD2 deletion and therefore SAC inactivation in HCT116 cells (Wild et al. 2018). Interestingly, truncation mutations in the APC/C subunit CDC27 are frequently found in cancer and cell lines engineered with these mutations display elongated mitotic timing and reduction of CIN (Sansregret et al. 2017). In agreement with these previous results, heterozygous deletions of either p31^comet^ or CDC16 in dipHAP1 cells showed resistance to SAC inhibition, extended mitotic timing, and were sufficient to reduce CIN. This indicates that the copy number changes of these genes were the drivers of the acquisition of the monosomies of chromosome arms 13q and 6p in the adapted cell lines. This delay in anaphase allows the chromosomes more time to biorient, which could potentially also suppress other sources of CIN (Thompson, Bakhoum, and Compton 2010). In general, our results demonstrate that specific aneuploidies can have impactful effects on cell cycle timing. How widespread this is in cancer and whether other stages of the cell cycle are also affected by specific aneuploidies is currently unknown.

Attempts to recapitulate cancer aneuploidy phenotypes in human cells and determine the genetic basis behind them have been extremely challenging (Ben-David and Amon 2019). Only recently there have been a few breakthroughs in identifying and understanding the role of individual frequent aneuploidies in human cells. For instance, resistance to the microtubule-stabilizing drug paclitaxel has been demonstrated via the acquisition of the loss of chromosome 10 in RPE1 cells (Lukow et al. 2021). In addition, a series of experiments using mouse and human cell models for Ewing sarcoma associated the gain of chromosome 8 with the upregulation of the Rad21 and Myc genes that reside on that chromosome (Su et al. 2021). Despite the challenges in determining the basis behind aneuploidy selection in human cells, there are multiple examples from studying aneuploidy in yeast (Selmecki, Forche, and Berman 2006; Rancati et al. 2008; Ravichandran et al. 2018; Chen et al. 2012; 2015). In these cases, the benefits of aneuploidy under selective conditions can often be attributed to only one or two genes. Whether this also holds true for the much larger human chromosomes was unknown.

Here, we identified two aneuploidies that were sufficient for a relative increase in resistance to the drug reversine and determined that a single gene on each chromosome is sufficient to recapitulate the phenotype when heterozygously deleted, both qualitatively and quantitatively. Our results suggest that single genes can provide the selective advantage behind the misregulation of hundreds of genes on a human chromosome. This result contrasts with computational models of aneuploidy patterns in cancer that attribute multiple oncogenes and tumor suppressors across the length of a chromosome for the selection of chromosome gains and losses (Davoli et al. 2013; Sack et al. 2018). Yet these differences between *in silico* models and *in vivo* models could be attributed to the differences in the diverse selective forces at play during tumorigenesis and the strong, specific selection from a small molecule inhibitor. Determining the basis behind the gains of chromosomes 5, 8 and 20 that are both common in cancer and observed in our adaptation experiments will help resolve this discrepancy. Furthermore, the contribution of single genes to aneuploidy selection could be more applicable to other processes such as chemotherapy resistance.

### Materials & Methods

#### Cell lines and cell culture

All cell lines used in this study have been tested negative for mycoplasma contamination. Their tissue types, sources, authentication and culture conditions are summarized in Supplementary Table 1. All cell lines were cultured in a humidified growth chamber at 37°C and 5 % CO2 in the respective media. The parental hTERT-RPE1 cell lines with inducible Cas9 (wildtype; inducible single knockout p53^-/-^) were kindly provided by I.M. Cheeseman (University of California). The parental HCT116 cell lines (wildtype and p53^-/-^) were gifts from B. Vogelstein (The Johns Hopkins Oncology Center). DLD1 was provided by M. Baccarini (Max Perutz Labs Vienna). The parental hTERT-HME1 cell line was obtained from Evercyte (Cat#: CHT-044-0236), the parental EEB is a cellosaurus cell line (Riken, RCB2345).

#### Generation of knockout cell lines

TP53 knockout: To generate the RPE1 TP53 single knockout, Cas9 was induced by adding 1µg/ml doxycycline hyclate (Sigma) every 24 hrs for 3 days before single-cell sorting. To generate the TP53 single knockouts in HME1, EEB and DLD1, a CRISPR/Cas9 was provided on a plasmid (Ran et al. 2013). SgRNA against Exon 2 of the TP53 gene was cloned into pSpCas9(BB)-2A-GFP (PX458, a gift from Feng Zhang, Addgene plasmid# 48138). The sgRNA plasmid was transfected into cells using FuGENE HD (Promega). 2 days after transfection, cells were sorted for the presence of Cas9 (GFP positive) and another 3 days later single cell sorting into 96-well plates was performed. Clonal populations were expanded gradually over the course of three weeks. TP53 mutations were identified by Sanger Sequencing and karyotypes of the cell lines were validated by whole-genome sequencing. p21, p31^comet^ and CDC16 knockouts: The knockouts in dipHAP1 and RPE1 cells were generated similar to the protocol above only XtremeGene 9 (Roche) was used as the transfection reagent for dipHAP1 cells and electroporation for RPE1 cells. All sgRNAs used for the generation of knockout cell lines are listed in Supplementary Table 5. The respective genotyping primers are listed in Supplementary Table 6.

#### Reversine adaptation assay

To initiate the adaptation process, parental p53^-/-^ cell lines were split into 12 independent populations at day 0 and reversine (Axon Medchem BV) was added at the indicated concentration for the respective cell line where ∼90 % of cells die (Extended Data Fig. 1a). 3 days later, single cell sorting into 96-well plates was performed and individualized cells were allowed to grow into colonies in the presence of reversine for the next 10-14 days. Around 20 independent cell populations for each cell line were expanded and continuously cultivated in medium containing reversine for up to 90 days. During this time period, the drug was replenished every 3-4 days. Reversine adapted populations were analysed at three time points, after 30, 60 and 90 days. For the last 30 days, reversine concentrations were increased to levels that are lethal for unadapted cells to further increase the selection of optimized adapted populations at 90 days (Extended Data Fig. 1b).

#### Next-generation sequencing (NGS) and data analysis

Genomic DNA from subconfluent populations was isolated using the QIAamp UCP DNA Micro Kit (Quiagen). NGS sample preparation was then performed as described earlier in Ravichandran et al., 2018. In brief, DNA samples were sheared to ∼500 base pairs (bp) fragments using the Bioruptor Pico sonicator (Diagenode) for two to three cycles (30 sec on/off). Fragmentation efficiency was monitored on a 0.8 % agarose gel stained with 1µM SYTOX green (ThermoFisher). DNA libraries were prepared with the NEBNext Ultra II DNA library kit for Illumina (NEB). DNA fragments with the optimal size were selected using AMPure XP beads (Beckman Coulter). Up to 96 cell lines per run were barcoded and multiplexed with NEBNext Multiplex Oligos (Index Primers 96well format, NEB) and mixed at equimolar ratios. The multiplexed samples were sequenced using the Illumina HiSeqV4 SR50 setting on an Illuina HiSeq 2500 system and using the Illumina NextSeq2000 P2 SR100 setting on an Illumina NextSeq 2000 system at the Vienna Biocenter Next-Generation Sequencing Facility (VBCF). All sequencing files are available on the Sequence Read Archive (BioProject ID: PRJNA885752). The demultiplexed data sets were then aligned to the human genome (assembly: GRCh38.p12) using Bowtie2 (version 2.2.9, http://bowtie-bio.sourceforge.net/bowtie2, (Langmead and Salzberg 2012)), converted to bed files using SAMtools (version 1.3.1; http://samtools.sourceforge.net, (Li et al. 2009; Li 2011)) and Bedtools (version 2.14, http://bedtools.readthedocs.io, (Quinlan and Hall 2010)). The resulting bed files were processed through the Gingko cloud software (Garvin et al. 2015) to correct for GC bias in low-read datasets and then analyzed with custom-made Python scripts.

#### In depth analysis of chromosome copy number changes

In depth characterization of chromosome copy number changes was carried out in 3 steps.

1. Segmentation and change point detection.
2. Assignment of segments to corresponding wild-type segments and determination of change state.
3. Correction of over-segmentation.

First, segmentation and change point detection was performed on copy number data that has been GC-debiased before using the Gingko online tool (Garvin et al., 2015). The array of binned copy number data points (each point represents 500 kb) is first split into segments of equal mean copy numbers that do not contain change points (locations where the mean copy number abruptly changes). By definition, this can be a whole-chromosome (no change point), a chromosome arm (one change point at the centromere, none within an arm) or a focal segment (at least one change point within a chromosome arm). The algorithm used to detect change points is PELT (Pruned Exact Linear Time) (Killick, Fearnhead, and Eckley 2012). By convention, if the algorithm finds a change point within a chromosome, it is split additionally at the centromere, to distinguish focal from arm copy number changes. Further, for acrocentric chromosomes only the data of q-arms was processed. The output after change point detection is a sorted list of segments.

For accurate determination of the change state of a segment, a 1-to-1 assignment between a sample and its corresponding wild-type cell line was implemented. For this, the difference between the mean copy number values of the sample segments and the wild-type segments was measured. Changes of the mean copy number value between sample and wild-type larger than 0.5 are marked as gain, changes smaller than -0.5 as loss. By convention, changed segments need to be at least 15 data points (=7.5Mb) long to get counted. If a change point is found in either the sample but not the wild-type or vice versa, an additional change point to ensure the 1-to-1 correspondence was introduced. For analyses that ignored pre-existing CNAs, certain segments were marked in the wild-type cell lines and excluded from further analysis. If a segment was flagged, all corresponding segments in the sample cell lines were also automatically flagged. The output of this analysis step is a list of segments, labeled with the change state (E=equal or no change, G=gain, L=loss). Flagged segments were labeled with the state i (=ignore). The alignment step described above has the disadvantage of over-segmentation of chromosome data, since it copies change points from wild-type data to the sample data (and vice versa) if the change point is present only in the wild-type. A second source of over-segmentation is the heuristic to split automatically at the centromere, if at least one change point is found. For correction of this over-segmentation sub-segments of the same change state were merged.

#### Colony formation assay

On day zero, 600 cells were seeded into 6-well plates and incubated for 12 days in their respective culture medium supplemented with reversine at the indicated concentrations (+Rev) or with the corresponding amount of DMSO (-Rev). The following reversine concentrations were used for the adapted cell lines: dipHAP1 400 nM; DLD1 50 nM; HCT116 100 nM; RPE1 125 nM; HME1 100 nM (Extended Data Fig. 1a). For the engineered dipHAP1 cell lines 300 nM reversine was used and for the engineered RPE1 cell lines 125 nM. Colonies were fixed with 4 % (vol/vol) formaldehyde in PBS (Thermo Fisher Scienitific) for 20 min, washed with purified water or PBS, stained for 20-30 min with Crystal Violet, washed with purified water several times and dried. The plates were imaged using a ChemiDocMP Imaging System (Bio-Rad) and analysis was done in ImageJ. The images were thresholded, and the fraction of area above the threshold was measured.

#### SiR-DNA staining & live cell imaging

48h before imaging, cells were seeded in 6- or 12-well glass bottom plates (Cellvis). On the day of imaging, cells were stained with 125 nM SiR-DNA (Spirochrome) in culture medium for 3h and afterwards maintained in Opti-MEM supplemented with 12.5 nM SiR-DNA and the indicated drugs during microscopy. For few cell lines (explicitly labeled in the figure legends), 10 µM Verapamil (Spirochrome, from SiR-DNA kit) was added to the staining and imaging medium to increase SiR-DNA incorporation. Automated microscopy of dividing human cells was performed on a Celldiscoverer 7 screening microscope (ZEISS), using a 50x water immersion objective (ZEISS Plan-APOCHROMAT 50x/ 1,2 W Autocorr and Autoimmersion Objective) with 0.5x zoom lens and recorded as three-dimensional time series. Each time series was 2.5 h long with images taken every 3 min. Imaging was controlled by ZEN 2.5 (blue edition) software. Cells were maintained at 37°C in a humidified atmosphere of 5 % CO2 throughout the entire experiment.

#### RNAi

The siRNA (Dharmacon) used in this study is listed in Supplementary Table 7. A non-targeting scrambled siRNA was used as a negative control. SiRNA transfections were performed using Lipofectamine RNAiMAX (ThermoFisher) according to manufactureŕs instructions and at room temperature. Diploid HAP1 and RPE1 cells were plated at 25-30 % confluence 12 hours prior to siRNA transfection in 6-well glass bottom plates. 9µl RNAiMAX was diluted in 150 µl OptiMEM (Gibco), 3 µl of 10 µM siRNA (30 pmol) was dissolved in 150 µl OptiMEM. Both solutions were combined, mixed by pipetting, and incubated for 5 min. The siRNA-lipid complex was then added dropwise to 6-wells and cells were incubated for 24 hours in the presence of the siRNA. Cells were subsequently subjected to SiR-DNA staining and live cell imaging as described above. Nocodazole was used at 250 ng/ml as a readout for siRNA efficacy.

#### DNA content analysis by flow cytometry

Human cells were trypsinized and stained with 0.2 µg ml^-1^ Hoechst 33342 (Thermo Fisher Scientific) for 30 min at 37°C. Analysis by flow cytometry immediately after incubation was done on a BD FACSAria^TM^ IIu Cell sorter (BD) using an argon laser tuned for UV (353-365nm) and fluorescence detection at 480 nm.

#### Fluorescence activated cell sorting (FACS)

FACS was performed on a BD FACSAria^TM^ IIu Cell sorter (BD) or on a BD FACSMelody^TM^ cell sorter (BD). For all cell lines and FACS experiments medium containing PBS and 10 % FBS was used. FACS was performed either in bulk or single cell sort into 96-well plates depending on the experiment. Growth medium for FACS sorted cells contained Normocin (Fisher Scientific) following the first two days after sorting to suppress mycoplasma, bacterial and fungal contamination.

#### p53 function assay and immunostaining

p53^+/+^ and p53^-/-^ cell lines were treated with 6.22 Gy γ-irradiation (Co60-Dmax), collected and fixed in ice-cold ethanol for at least 20 min. After ethanol removal, fixed cells were washed twice with PBS and then permeabilized with 0.25 % Triton X-100 in PBS for 10 min at 4°C. Cells were blocked for 1h with 1 % BSA in PBS and stained for 2 hours with a mouse monoclonal MPM-2 Antibody (10 µg/ml; Abcam). After several washes, cells were stained with Alexa 488-conjugated goat anti-mouse secondary antibody (10 µg/ml; Thermo Fisher Scientific). DNA was stained with Propidium Iodide-Rnase solution (PI, final conc.: 50 µg/ml + 100 µg/ml RNAse A in PBS) in the dark at RT for another 20 min and then subjected to FACS analysis.

#### Engineering of full and partial chromosome arm deletions

For the engineering of chromosome arm deletions the CRISPR/Cas9 system (Ran et al. 2013) was used based on an adapted approach from Zuo et al. (2017). Guide RNAs that target repetitive sequences within the chromosome arm were selected based on a custom-made Python script. Additionally, single cutting sgRNAs targeting the intergenic region either close to the centromere or at the respective partial deletion sites were selected based on their top ranking in the online tool GuideScan (https://guidescan.com, (Perez et al. 2017), homepage recently updated to GuideScan2 (Schmidt et al. 2022)) and CRISPOR (http://crispor.tefor.net, (Concordet and Haeussler 2018)). Off-target analysis was performed using Cas-OFFinder (CRISPR RGEN Tools, http://www.rgenome.net/cas-offinder/, (Bae, Park, and Kim 2014)). All sgRNAs used for engineering of whole or partial chromosome deletions are listed in Supplementary Table 3 and Extended Data Fig. 7b. The number of protein coding genes was calculated based on the MANE project (release v1.0, (Morales et al. 2022)).

For full and partial chromosome arm deletions, dipHAP1 p53^-/-^ cells were seeded subconfluently in a 6-well plate one day before transfection. 1µg of pSpCas9(BB)-2A-GFP plasmid was transfected with XtremeGene9 (Roche). 48 h after transfection, cells were FACS sorted for GFP expression either in bulk or as single cells in 96-well plates. Three days later, bulk sorted cells were single cell sorted in 96-well plates. Cell clones were expanded to confluency in a 12-well plate. Genomic DNA was extracted with 0.5x lysis buffer (Ramlee et al. 2015) and DNA concentration was measured with the Qubit dsDNA HS assay (Invitrogen) according to manufacturer’s protocol. The generation of aneuploidy was ascertained by quantitative PCR (qPCR) according to a modified protocol from Ravichandran et al. 2018. In brief, Luna universal qPCR master mix (NEB) and 10 ng DNA of each sample were used together with two primer pairs targeting the telomeric and the centromeric region of the respective chromosome arm (Extended Data Fig. 7b, Supplementary Table 4). All qPCR reactions were performed on Mastercycler ep realplex platforms (Eppendorf). Each calculated cycle threshold (Ct) value was normalized to the Ct value generated by a primer pair targeting the euploid chromosome 4 (ALB control primer). Samples and controls were measured in triplicates. Additional to qPCR analysis, the karyotype of all engineered aneuploid cell lines was verified by whole genome sequencing according to the above-described protocol. The aneuploid cell lines were frozen in multiple aliquots and for each experiment fresh aliquots were thawed and cultivated for a maximum of 6 passages to reduce the formation of secondary karyotypic changes.

#### Co-cultivation assay

GFP-expressing and non-fluorescent (dark) cell lines were seeded separately in 10 cm dishes prior to FACS sorting. The two cell lines were then mixed in a 50:50 ratio by bulk sorting 400,000 cells each into 4 ml of growth media with 1:500 Normocin. Setting up the gates was done using a non-fluorescent (dark) parental cell line and a GFP expressing cell line as controls (Extended Data Fig. 10b). Mixed cell lines were cultured in standard growth media or growth medium supplemented with 300 nM reversine in six replicates for each growth condition, were 50,000 of mixed cells were added into each well of a 6-well plate. The remaining mixed cells were used to determine the exact distribution of GFP and dark cells at day 0 by flow cytometry. After 4 days, cell mixtures were passaged, and the relative abundance of GFP was measured by flow cytometry. All later measurements were done in intervals of three to four days for a total duration of 16 - 19 days.

#### Western blotting

For protein preparation 2 or 4 million cells were lysed in 75 µl ice cold NP40-buffer with cOmplete protease inhibitor cocktail (Roche) and incubated on ice for 1h. Samples were then incubated with 0.1 µl Benzonase nuclease (Sigma) for 30 min at 37°C. 25 µl 4x protein sample buffer was added and boiled at 95°C for 5 min. Protein size was estimated using the pre-stained protein Ladder – mid range molecular weight (Abcam). Coomassie staining was used to equalize the amount of each sample for loading. SDS-Page was performed at 120 V for 90 min. Wet transfer onto a nitrocellulose membrane was performed at 250 mA for 150 min at 4°C. Membranes were blocked for 1 h in 5 % BSA (Sigma) or 5 % skim-milk (Gerbu Biotechnik GmbH) in Tris-buffered saline with 0.01 % Tween (TBST), depending on the primary antibody (see below). Primary antibodies diluted in the respective blocking buffer were incubated overnight at 4°C on an orbital shaker. The membranes were washed 30 min in TBST with three buffer exchanges before and after secondary antibody incubation. Horseradish peroxidase (HRP) conjugated anti-mouse IgG secondary antibody (1:10.000, Cat. #7076; Cell Signaling Technology) diluted in the respective blocking buffer was incubate for 1h at room temperature. Chemiluminescence was developed using the Amersham ECL Prime detection reagent (Cytiva) and detected with the ChemiDocMP Imaging System (Bio-Rad). Quantification of protein bands was carried out using ImageJ. The following primary antibodies were used: anti-CDC16 (in skim-milk-TBST, 1:500, mouse, catalogue no. (E-4):sc-365636; Santa Cruz Biotechnology), anti-p31^comet^ (in BSA-TBST, 1:500, mouse, catalogue no. E29.19.14; Sigma-Aldrich), anti-alpha-Tubulin (in BSA and skim-milk-TBST respectively, 1:20,000, mouse, catalogue no. T6074; Sigma-Aldrich).

#### Statistical analysis and sample number

All experiments were repeated multiple times and the indicated number of experiments always refer to biological replicates. Statistical analysis was performed using Python 3.4 and GraphPad Prism software. For statistical analysis of differences to the wild type cell lines, the unpaired student’s t-test was used. Details for the statistical tests used in a particular experiment are reported in the figure legends. Error bars represent +/-standard deviation (SD) unless otherwise indicated.

## Acknowledgements

The authors thank the Campbell and Dammermann laboratories for helpful discussions and comments. We thank the Loizou, Vogelstein, Cheeseman, Baccarini, and Versteeg labs for gifts of cell lines and plasmids. We thank David Teis, Dea Slade, and Alex Dammermann for careful reading of the manuscript. We acknowledge the service of the Max Perutz Labs BioOptics Light Microscopy and FACS Facilities, members of the VLSI. We would also like to acknowledge the services provided to us by the VBC sequencing facility located at the VBCF.

This work was supported by Vienna Science and Technology Fund (WWTF) grant VRG14-001 and Austrian Science Fund (FWF) grants Y944-B28 and W1238-B20 to C.S.C.

## Author Contributions

M. A. Y. A. and T. C. K. conducted the majority of the experiments. R. H. generated a phyton based pipeline for high resolution CNV detection in NGS data. M. A. Y. A. performed the NGS analysis. L. W. performed all Western blot experiments. M. A. Y. A. and J. S. generated most of the gene knockout cell lines. A. Mark. performed the co-cultivation experiments. A. Mart. screened for mutations in MPS1. M. A. Y. A., T. C. K. and C. S. C conceptualized the research, analyzed the data, and wrote the manuscript. All authors reviewed and approved the manuscript.

---

**Extended Data Figure 1.**
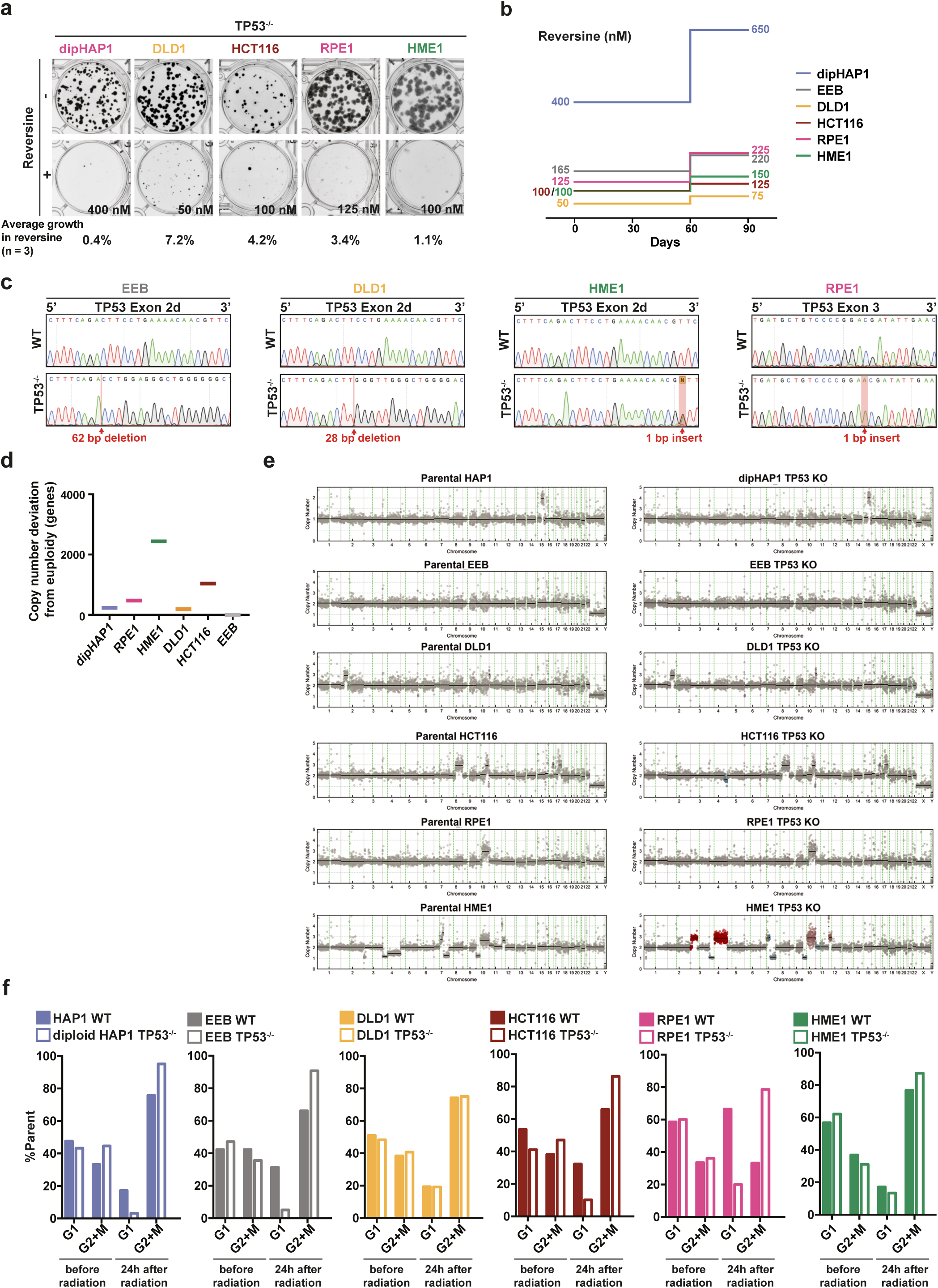
**a**, Growth of p53 knockout cell lines in the presence or absence of reversine in colony formation assays. Representative images and averaged growth from three independent experiments are shown. **b**, Reversine concentrations that were used during adaptation of the respective cell lines. **c**, DNA sequencing chromatograms of the TP53 target sites of wildtype and the CRISPR/Cas9 mutated cell lines. **d**, Copy number deviations from euploidy in p53 knockout cell lines before adaptation as measured by next generation sequencing. Copy number changes are depicted as number of genes. **e**, Copy number profiles of wildtype cell lines and corresponding p53 knockouts. Shown are copy numbers with each dot representing a 500 kb bin in the genome of cell populations. Segments above 0.5 threshold for gain are colored in red, segments below threshold for loss are colored in blue. Thresholding occurs relative to the parental wildtype. Dotted green lines mark the centromeres in each chromosome. **f**, Bar graphs showing flow cytometric analysis of wildtype (full bar) and corresponding p53 knockout (empty bar) cell lines before and 24 hours after radiation with 6.22-Gy. Cells were stained with PI and anti-MPM-2. Flow cytometry was done as described in the methods. 10,000 cells were analyzed in each experiment. S-Phase percentages are not shown.

**Extended Data Figure 2.**
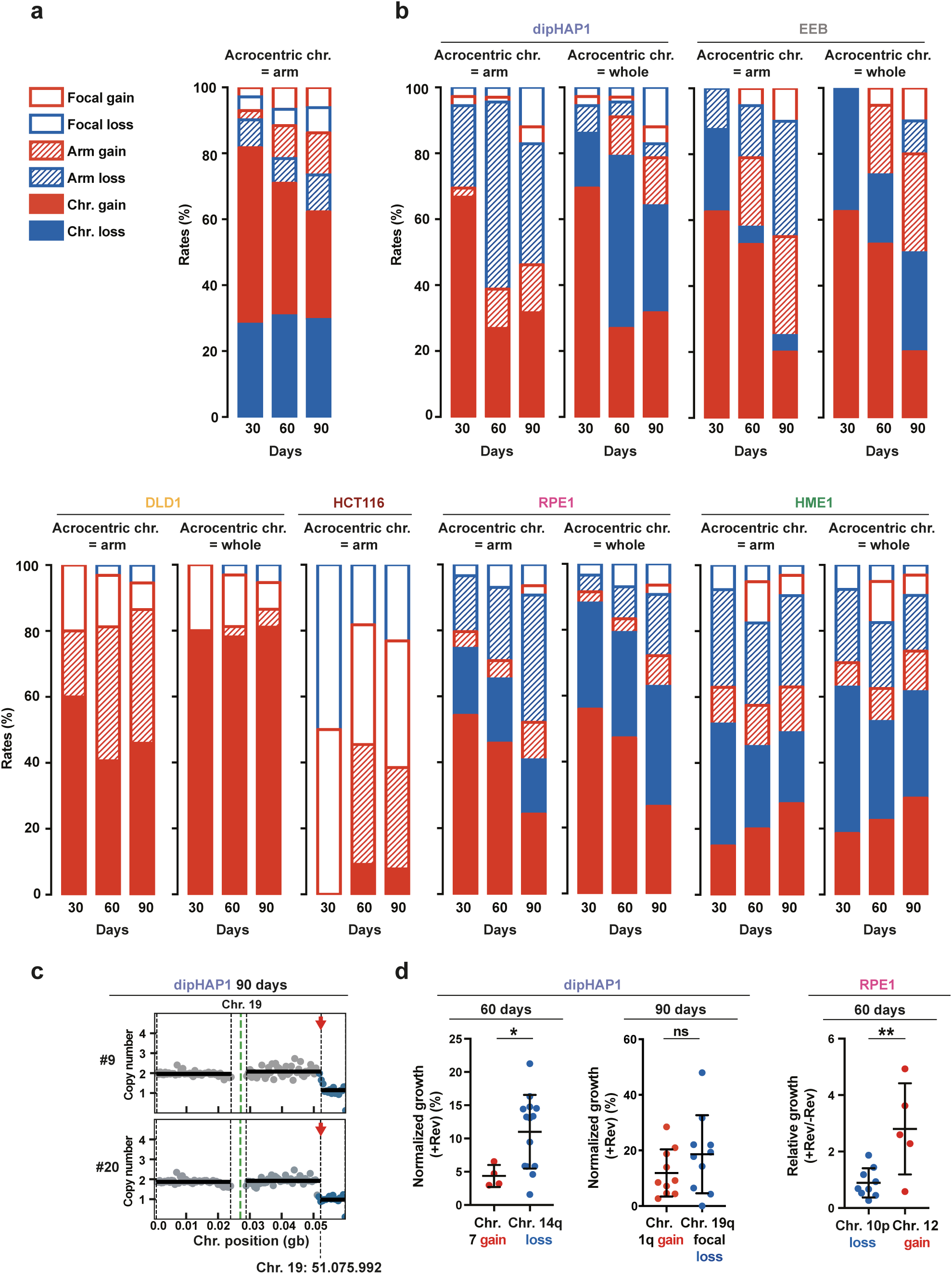
**a**, Relative proportion of types of copy number changes as in Fig. 1e with acrocentric chromosomes considered as whole-chromosomes instead of arms. **b**, Relative proportion of types of copy number changes of the reversine cultivated cell populations at 30, 60 and 90 days from each cell line. For acrocentric chromosomes considered as arm or as whole chromosome. **c**, Copy number profiles of two dipHAP1 cell lines (#9 and # 20) with a 7.5 MB focal loss of chr. 19q. **d**, Comparison of the normalized growth with reversine (+Rev) or relative growth (+Rev/-Rev) of 60 or 90 days adapted dipHAP1 and RPE1 populations with the negatively correlated aneuploidies related to Fig. 1g. p-values are from unpaired student’s t-test. * = p<0.05, ** = p<0.01, ns = not significant.

**Extended Data Figure 3.**
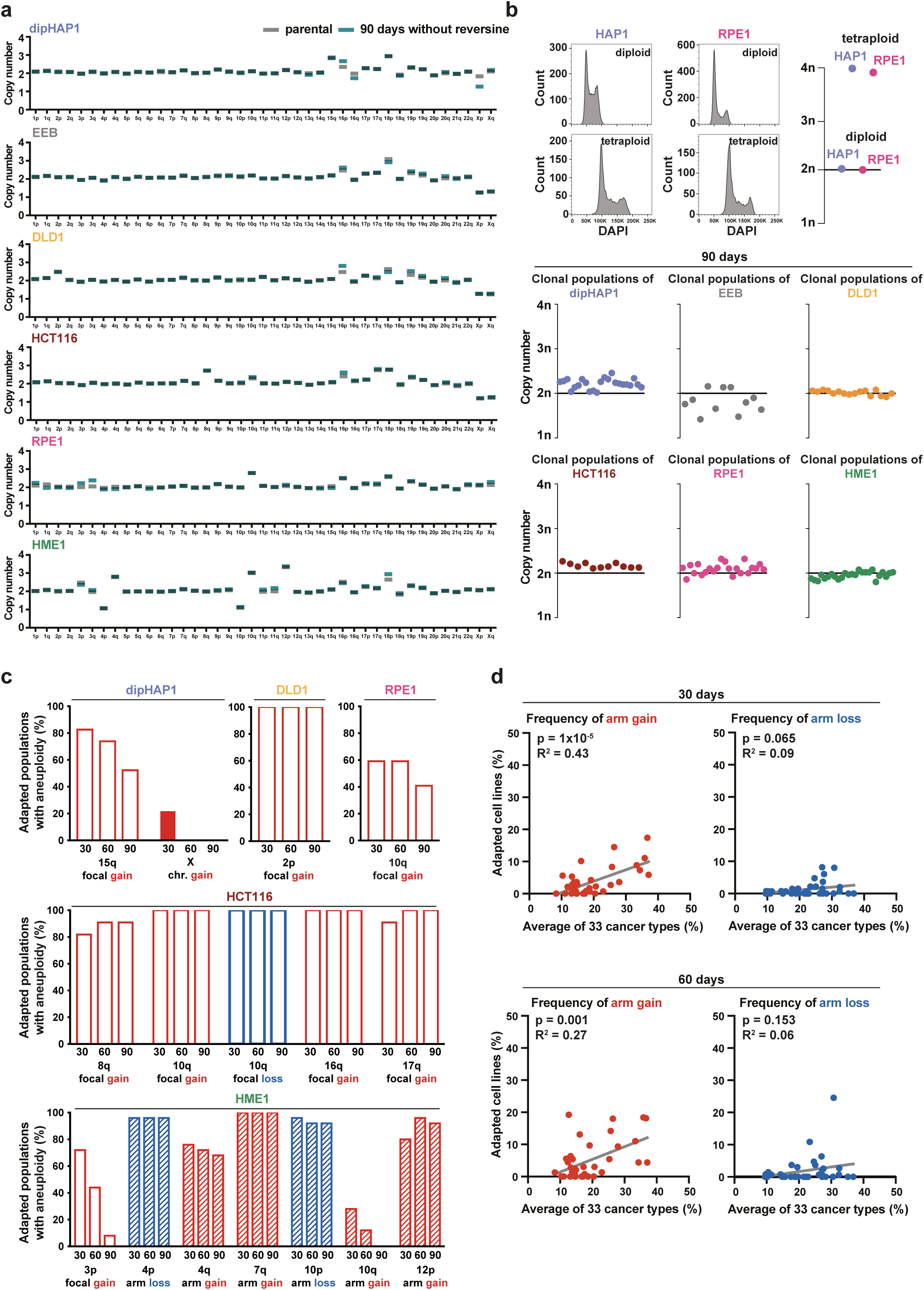
**a**, Genome-wide arm copy numbers of parental cell lines before adaptation (time point zero, grey) and after 90 days of cultivation in the absence of reversine (turquoise). 90 days values are an average of six independently cultivated populations. **b**, HOECHST-based flow cytometry analysis of DNA content of diploid and tetraploid dipHAP1 and RPE1 control cell lines (top). G1 DNA content of the reversine adapted populations of all six cell lines at the 90 days time point (bottom). 10,000 cells were analyzed in each measurement; 1n represents haploid DNA content. **c**, Bar graphs depicting the relative abundance of cell line-specific aneuploidies present at time point zero after 30, 60 and 90 days of reversine adaptation. Bar colors depict type of copy number change as in Fig. 1e. **d**, Correlations between the frequencies of chromosome arm gains (red dots) and losses (blue dots) of reversine adapted cell populations at the 30 days and 60 days time points and 33 different cancer types. The R-squared values and p-values from F-tests are depicted in the graphs.

**Extended Data Figure 4.**
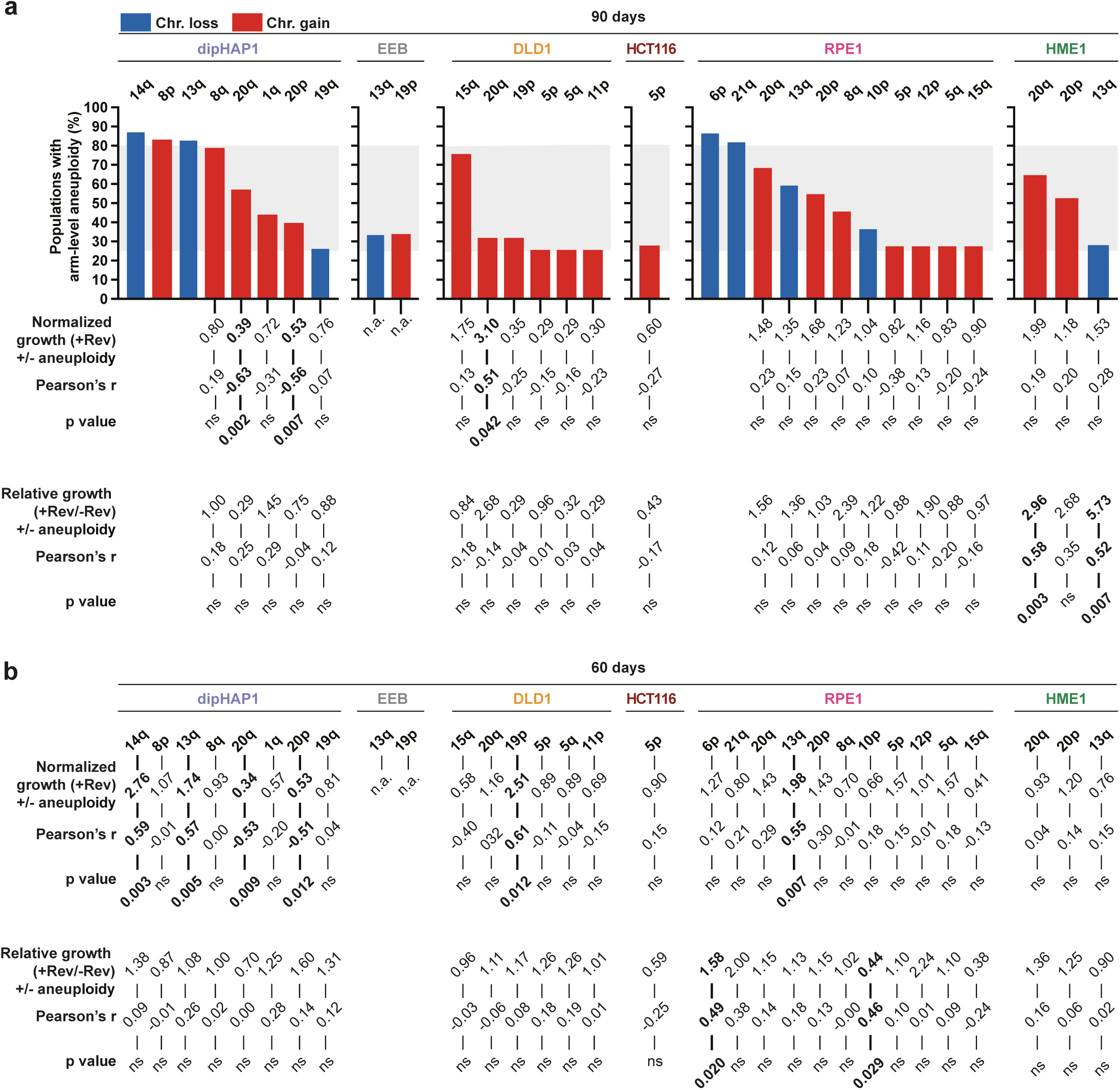
**a**, Bar graphs representing all arm aneuploidies that are present in >25% of reversine-adapted populations of a given cell type. The grey areas depicts the range (25-80%) in which correlation analysis between copy number changes and fitness was performed for the 90 days time point. Normalized growth with reversine (+Rev) and relative growth (+Rev/-Rev) of the adapted populations with and without the respective aneuploidy as well as corresponding correlation coefficients and p-values are depicted. Growth values were obtained from two independent experiments, **b,** Correlation analysis as in **a,** only for the 60 days time point, p-values are from unpaired student’s t-test. ns = not significant, n.a. = not assessed.

**Extended Data Figure 5.**
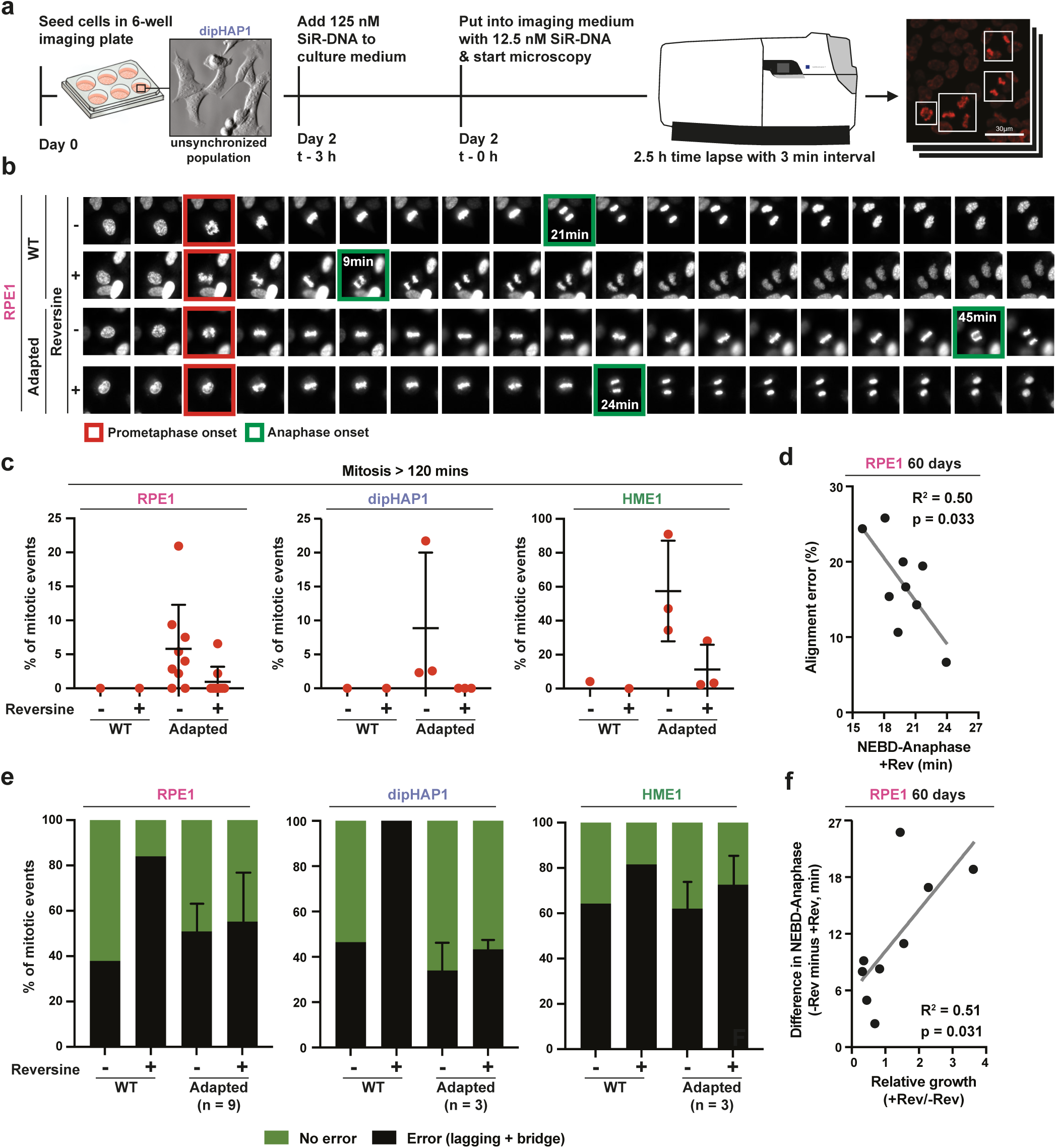
**a**, Scheme of the SiR-DNA staining procedure followed by live cell time-lapse imaging. **b**, Full mitotic time-lapse with and without reversine of the parental and adapted RPE1 cells shown in Fig. 4a. **c**, Percentage of mitotic events that lasted longer than 120 min, related to Fig. 4a,b. Each dot represents the mean percentage of an individual adapted population of the respective cell line or the respective parental cell line (WT). **d**, Correlation between the percentage of mitotic events with alignment defects directly preceding anaphase (y-axis) and the NEBD to anaphase duration in reversine (x-axis) of nine 60 days adapted RPE1 population, related to Fig. 4a,c. **e**, Quantification of anaphase errors (bridges and lagging chromosomes) with and without reversine for the unadapted parental cell lines and the adapted cell populations from RPE1 (time point 60 days), dipHAP1 (time point 90 days) and HME1 (time point 90 days). Error bars depict SD between the adapted populations (n depicted in graph) of the respective cell line. **f**, Correlation between the difference in NEBD to anaphase timing with and without reversine (y-axis) and the relative growth (+Rev/-Rev) of the nine 60 days adapted RPE1 populations (x-axis), related to Fig 4a. Analyzed adapted populations are the same as in Fig. 4a-d. p-values are from F-tests.

**Extended Data Figure 6.**
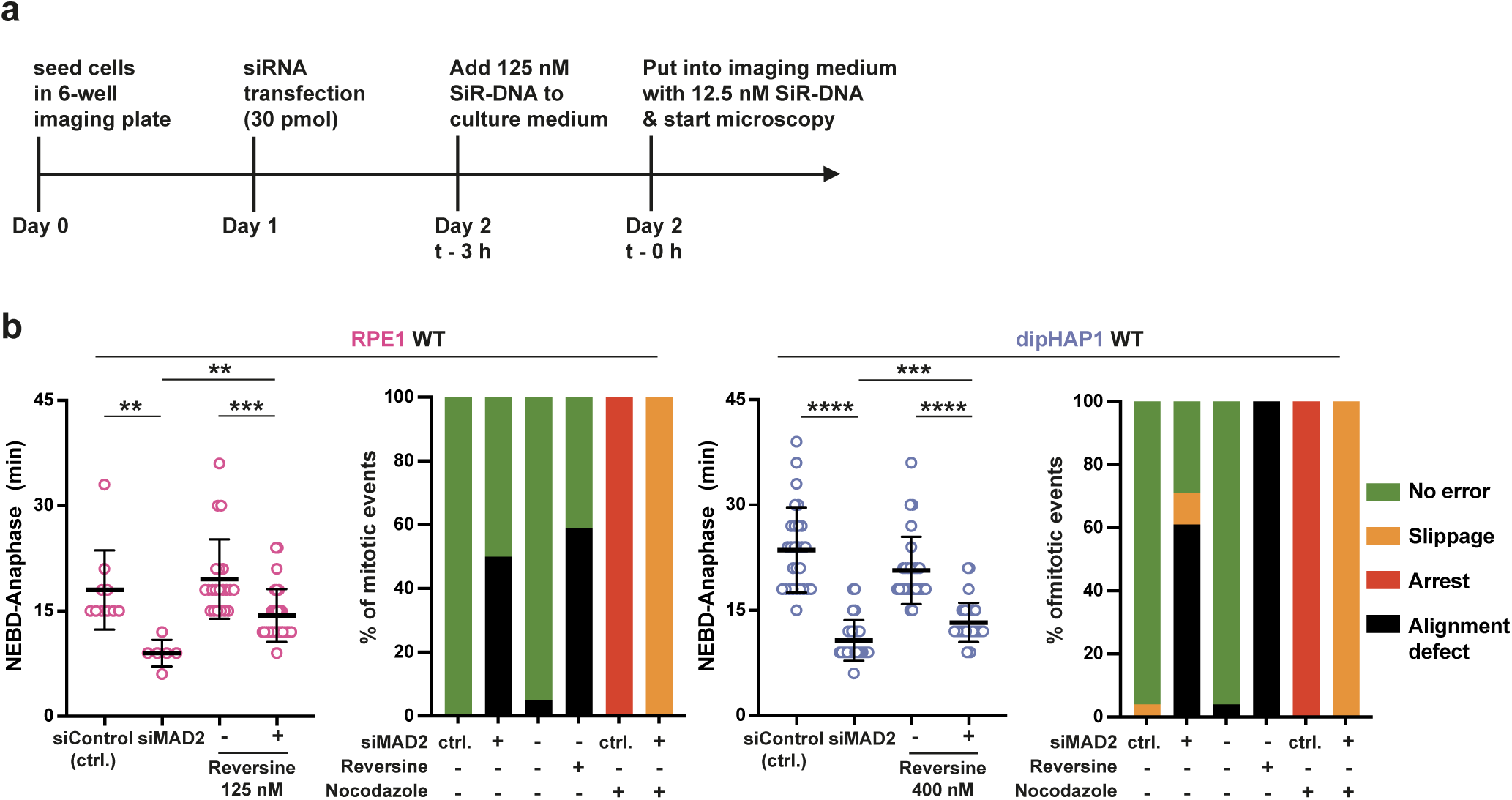
**a**, Experimental design of the siRNA experiment targeting MAD2 combined with the SiR-DNA live cell imaging set-up. **b,** Quantification of NEBD to anaphase durations and mitotic errors of RPE1 and dipHAPI unadapted parental (WT) cell lines. Left: Time from NEBD to anaphase in min of WT cell lines transfected either with the indicated siRNA or with and without reversine. Right: Quantification of mitotic errors under the indicated conditions. Green = no error, orange = slippage, red = mitotic arrest, black = alignment defect, p-values are from unpaired student’s t-tests, ** = p<0.01, *** = p<0.001, **** = p<0.0001.

**Extended Data Figure 7.**
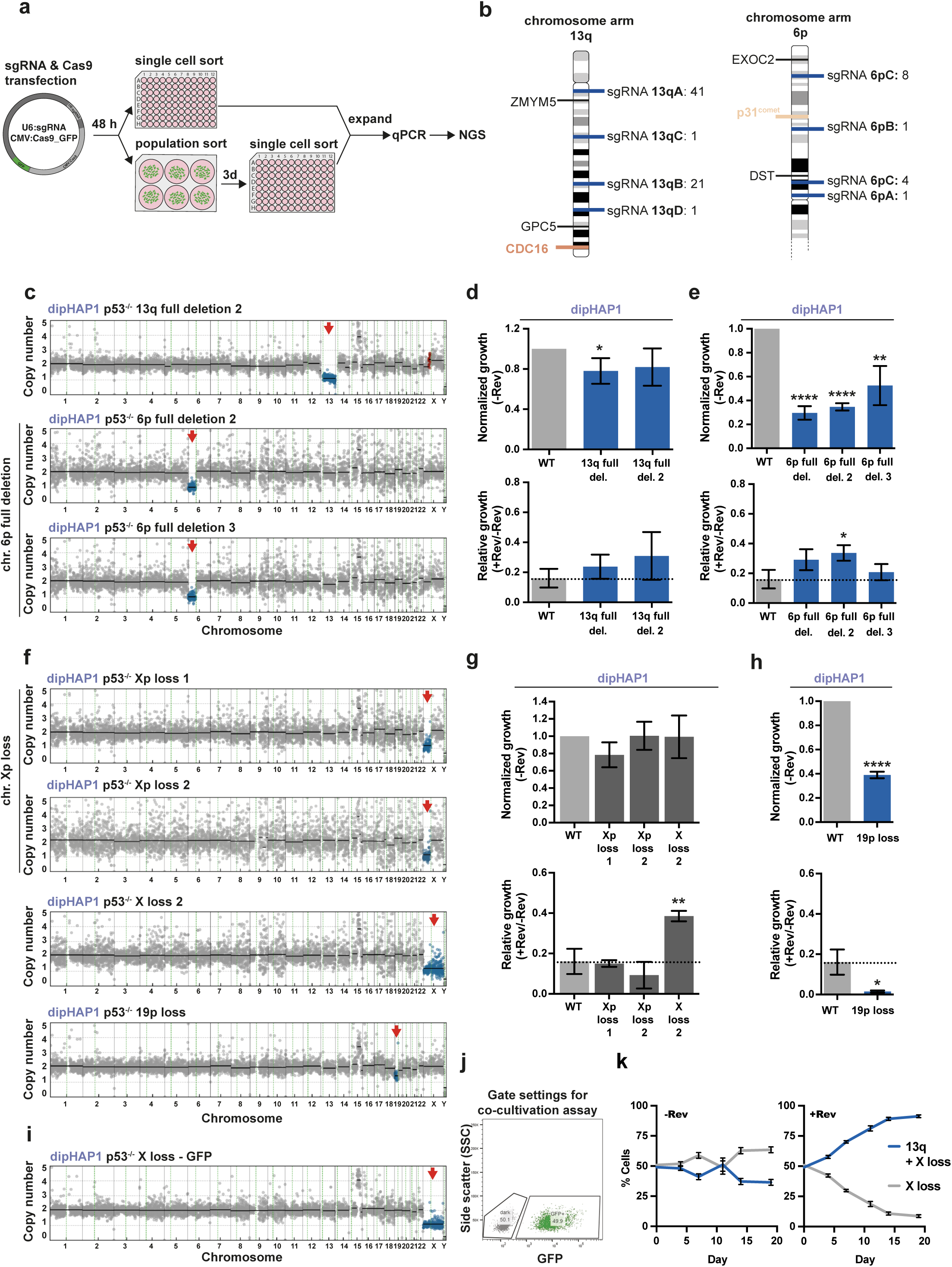
**a**, Illustration of the experimental procedure to engineer whole and partial chromosome deletions. **b**, Schematic depiction of sgRNA target sites on chromosome arm 13q and 6p (blue lines). Numbers after the semicolon denote the number of target sites on the chromosome arm. The qPCR primer binding locations that were used to screen for successful deletion cell lines are depicted with black lines and the gene name. **c**, Copy number profiles of additional engineered chromosome 13q (13q full del. 2) and 6p deletions (6p full del. 2, 3) in dipHAP1 cells. **d**,**e** Normalized and relative growth of the additional 13q and 6p partial deletion cell lines measured by colony formation assays. Bars depict mean +/-SD. Dotted line represents mean growth of dipHAP1 WT cell line in reversine. n = 3. **f,** Copy number profiles of the dipHAP1 Xp (Xp loss 1 and 2), additional X loss (X loss 2) and 19p loss cell lines. **g,h** Normalized and relative growth of the dipHAP1 Xp, X loss and 19p loss cell lines measured by colony formation assays. n = 3. **i**, Copy number profile of the X loss-GFP cell line that was used for the co-cultivation assay. **j**, Schematic representation of the gates used for the co-cultivation assay both for the initial cell sorting by FACS and the following measurements. **k**, Second independent co-cultivation of the 13q + X loss cell line and the X loss-GFP cell line. The mean +/-SD for 6 separate populations of cells in one time course is shown. p-values are from unpaired student’s t-tests, * = p<0.05, ** = p<0.01, **** = p<0.0001. Where not indicated, the p-value was greater than 0.05.

**Extended Data Figure 8.**
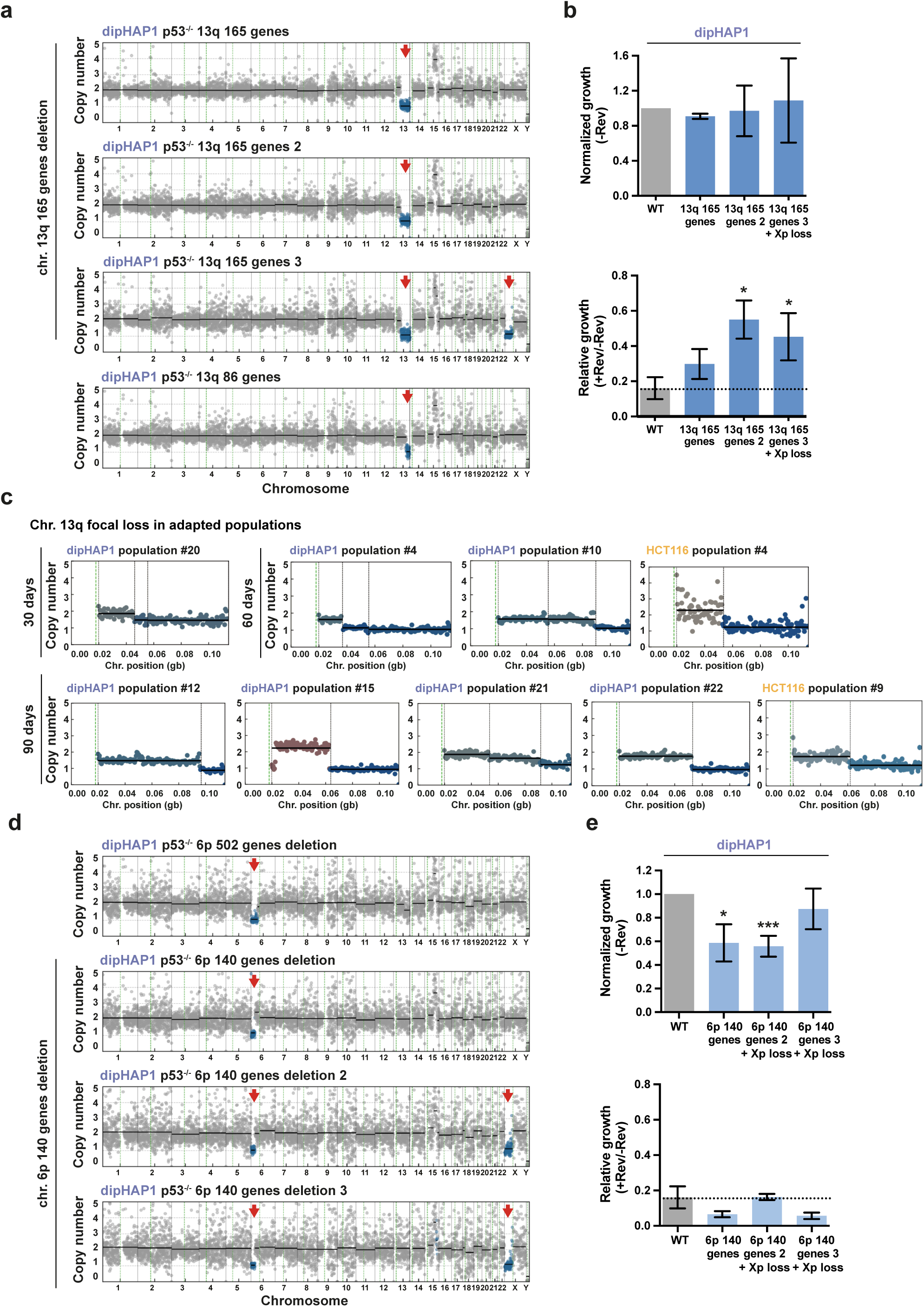
**a**, Copy number profiles of all engineered dipHAP1 chromosome 13q partial deletions. Cell line 13q 165 genes 3 also lost chromosome arm Xp. **b**, Normalized and relative growth of the additional 13q partial deletion cell lines (13q 165 genes 2 and 3). Bars depict mean +/-SD. Dotted line represents mean growth of dipHAP1 WT cell line in reversine. n = 3. **c**, Copy number profiles of 13q focal deletions that arose in adapted populations of the dipHAP1 and HCT116 cell lines after 30, 60 or 90 days of adaptation as indicated. **d**, Copy number profiles of all engineered dipHAP1 chromosome 6p partial deletions. Cell lines 6p 140 genes 2 and 3 also lost chromosome arm Xp. **e**, Normalized and relative growth of all 6p partial deletion cell lines. n = 3. p-values are from unpaired student’s t-tests, * = p<0.05, *** = p<0.001. Where not indicated, the p-value was greater than 0.05.

**Extended Data Figure 9.**
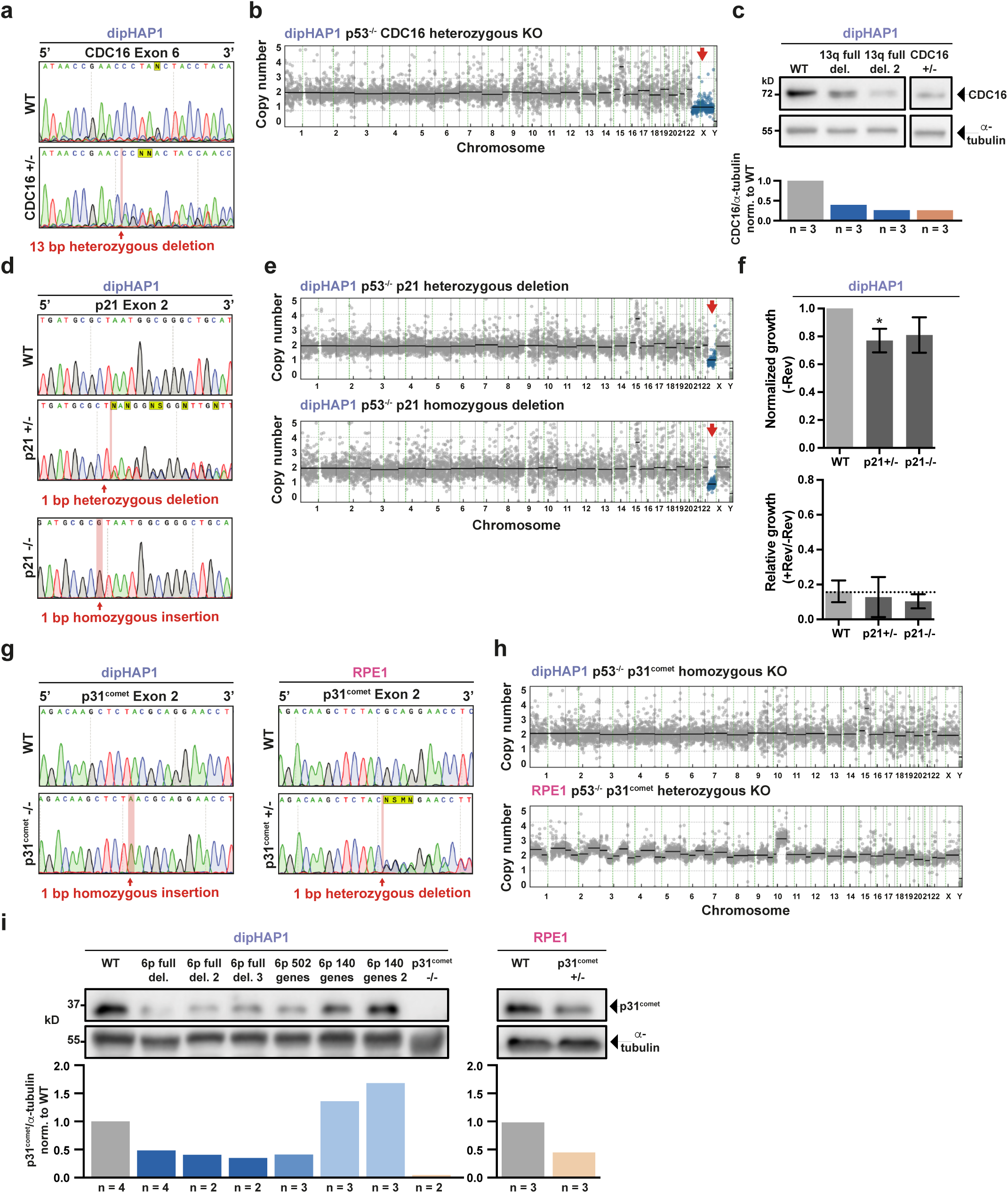
**a**, DNA sequencing chromatogram of the heterozygous CDC16 knockout in dipHAP1. **b**, Copy number profile of the dipHAP1 CDC16+/-knockout cell line with the additional monosomy of chromosome X. **c**, Western blot of CDC16 for the engineered chromosome 13q full deletion and CDC16+/-knockout cell lines. Alpha-tubulin was used as a loading control. The mean of the CDC16 intensity relative to the alpha-tubulin intensity and normalized to the WT is depicted in the bar graph; the respective n values are indicated below. **d,** DNA sequencing chromatograms of the heterozygous and homozygous p21 knockout in dipHAP1. **e**, Copy number profiles of the dipHAP1 p21+/- and p21-/-knockout cell lines with the additional loss of chr. Xp. **f**, Normalized and relative growth of the dipHAP1 p21+/- and p21-/-knockout cell lines. Bars depict mean +/-SD. Dotted line represents mean growth of dipHAP1 WT cell line with reversine. n = 2. p-values are from unpaired student t-tests, * = p<0.05. Where not indicated, the p-value was greater than 0.05. **g**, DNA sequencing chromatograms of the heterozygous and homozygous p31^comet^ knockout in dipHAP1 and RPE1. **h**, Copy number profiles of the heterozygous and homozygous p31^comet^ knockout cell lines. **i**, Western blot of p31^comet^ for all engineered chromosome 6p deletion and knockout cell lines. Alpha-tubulin was used as a loading control. Mean value of p31^comet^ intensity relative to alpha-tubulin and normalized to WT is depicted in the bar graph; the respective n values are indicated below.

**Extended Data Figure 10.**
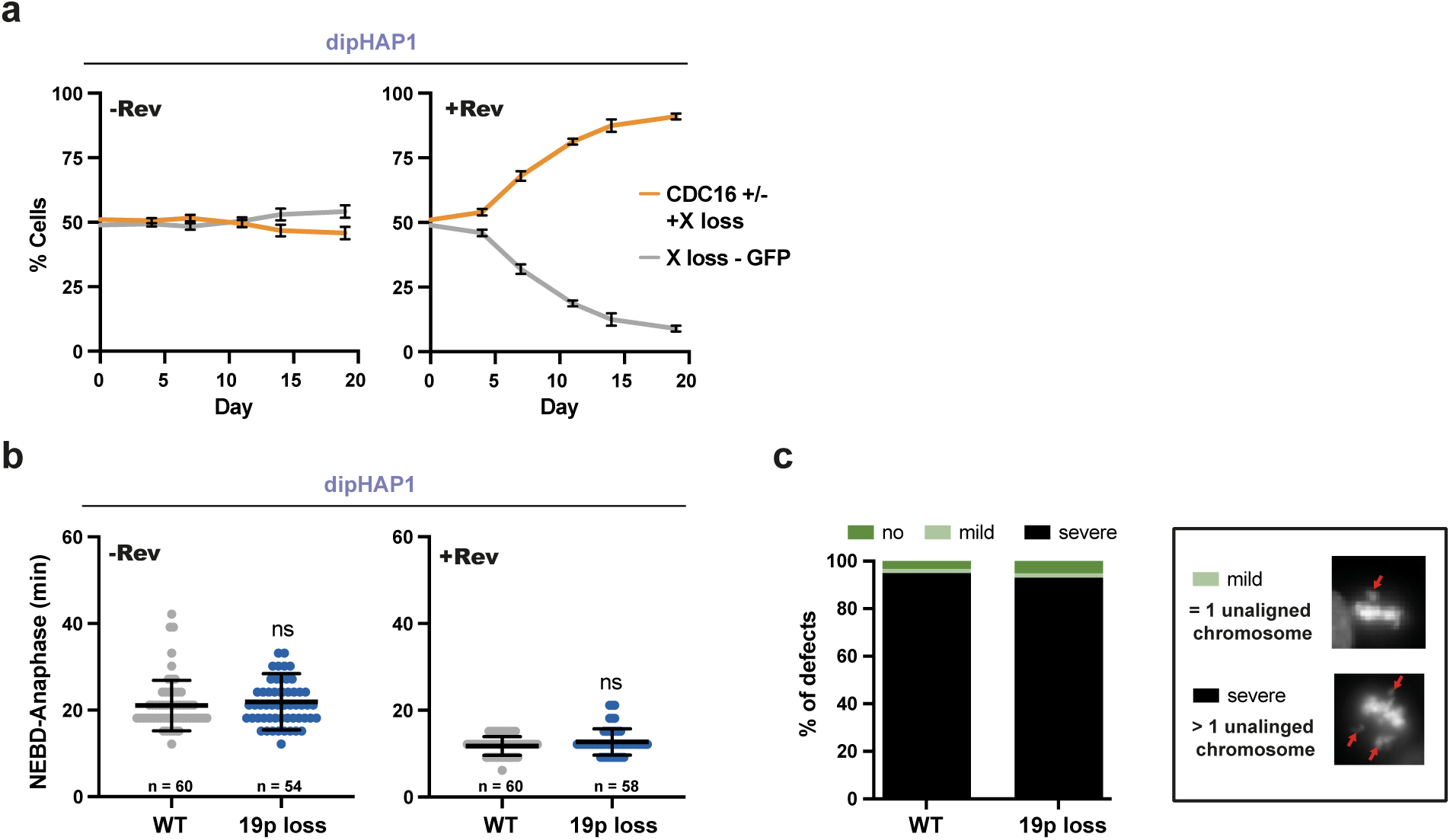
**a**, Second independent co-cultivation of the dipHAPI CDC16 +/-+ X loss cell line and the X loss-GFP cell line, related to Fig. 6c. The mean +/-SD for 6 separate populations of cells in one time course is shown, **b,** Duration from NEBD-anaphase in min for the dipHAPI 19p loss cell line without (-Rev), (n = 58-60 mitoses), and with reversine (+Rev), (n = 54­60 mitoses), n = 3. p-values are from unpaired student t-tests, ns = not significant, **c,** Quantification of no (dark green), mild (one unaligned chromosome, light green) or severe (multiple unaligned chromosomes, black) alignment defect of the analyzed mitoses from **b.** Example images for the categorization of mitoses into mild (one unaligned chromosome) or severe (multiple unaligned chromosomes) alignment defect.

