## Supplemental Tables for "Adaptation to spindle assembly checkpoint inhibition through the selection of specific aneuploidies"

| Cell line | Tissue | Phenotype | Sex | Culture Medium | Source |
|---|---|---|---|---|---|
| **hTERT-RPE1** | Retina epithelial | Adherent | Female | DMEM (Sigma-Aldrich)<br>+ 2 mmol L-Glutamine (Gibco)<br>+ 1mM Natrium-Pyruvate (Sigma-Aldrich) | I. M. Cheeseman (University of California, USA) |
| **hTERT-HME1** | Mammary gland epithelial | Adherent | Female | MEBM (Lonza)<br>+ 2 mmol L-Glutamine (Gibco)<br>+ 1mM Natrium-Pyruvate (Sigma-Aldrich) | Evercyte (Cat#: CHT-044-0236) |
| **DLD1** | Colorectal cancer | Adherent | Male | RPMI-1640 (ATCC) (Sigma-Aldrich) | M. Baccarini (Max Perutz Labs, Austria) |
| **HCT116** | Colorectal cancer | Adherent | Male | McCoy's 5A (Sigma-Aldrich)<br>+ 2 mmol L-Glutamine (Gibco)<br>+ 1mM Natrium-Pyruvate (Sigma-Aldrich) | B. Vogelstein (The Johns Hopkins Oncology Center, USA) |
| **diploid HAP1 (dipHAP1)** | Chronic myeloid leukemia (CML) | Adherent | Male | IMDM (Sigma-Aldrich) | J.I. Loizou (CeMM, Austria) |
| **EEB** | Acute myeloid leukemia (AML) | Suspension | Male | SFEM (STEMCELL Technologies) | Riken (RCB2345) |

All culture media were supplemented with 10% FBS and 1% Penicillin-Streptomycin.

**Supplementary Table 1.** Cell lines used in this study and their culture medium.

| Cell line | Karyotype deviations from euploidy |
|---|---|
| **parental RPE1** | Segmental chr. 10q gain |
| **RPE1 TP53 KO** | Segmental chr. 10q gain |
| **parental HME1** | segmental loss of 3p, loss of chr. 4, segmental gain & loss of chr. 7q, chr. 10 isochromosome, chr. 12p gain |
| **HME1 TP53 KO** | segmental gain of 3p, loss of chr. 4p, gain of chr. 4q, segmental gain & loss of chr. 7q, chr. 10 isochromosome, chr. 12p gain |
| **parental HAP1** | Segmental chr. 15q gain |
| **dipHAP1 TP53 KO** | Segmental chr. 15q gain |
| **parental EEB** | none |
| **EEB TP53 KO** | none |
| **parental DLD1** | Segmental chr. 2p gain |
| **DLD1 TP53 KO** | Segmental chr. 2p gain |
| **parental HCT116** | Segmental gains of chromosomes 8q, 10q 16q and 17q |
| **HCT116 TP53 KO** | Segmental gains of chromosomes 8q, 10q, 16q and 17q, segmental 4q loss (above 1.6 threshold) |

**Supplementary Table 2.** Karyotype deviations of the parental cell lines used in this study.

| Name | Sequence | Reference |
|---|---|---|
| **Chromosome arm 6p deletion** | | |
| **sgRNA 6p A** | ACGGTTTCATTAGTCATACC | This paper |
| **sgRNA 6p B** | GGGACCGTCACCCTAATAGG | This paper |
| **sgRNA 6p C** | TGGAATATTGTTCACCTTTA | This paper |
| **Chromosome arm 13q deletion** | | |
| **sgRNA 13q A** | GGGGGAGTGAATGTGAGTGA | This paper |
| **sgRNA 13q B** | ATATATGGGGTATACGTATA | This paper |
| **sgRNA 13q C** | TGGGTTACTTACCGACCGTG | This paper |
| **sgRNA 13q D** | GATAATACGATAGGCCAGTG | This paper |

**Supplementary Table 3.** sgRNAs that were used for the generation of whole or partial chromosome deletions. Further Illustration in Extended Data Figure 7b.

<table>
<tr><th>Name</th><th></th><th>Sequence</th><th colspan="2">Reference</th></tr>
<tr><td colspan="5">Chromosome 6p deletions</td></tr>
<tr><td rowspan="2">EXOC2</td><td>fw</td><td>ATGTCTCGATCACGACAACCC</td><td rowspan="2">PrimerBank, ID: 30581133c1</td><td rowspan="9">Spandidos et al. 2010</td></tr>
<tr><td>rev</td><td>GGCCAGTCCCCAGATTTTCT</td></tr>
<tr><td rowspan="2">DST</td><td>fw</td><td>CTACCAGCACTCGAACCAGTC</td><td rowspan="2">PrimerBank, ID: 291290967c1</td></tr>
<tr><td>rev</td><td>GCCGAAGCTAATGCAAGAGTTG</td></tr>
<tr><td colspan="4">Chromosome 13q deletions</td></tr>
<tr><td rowspan="2">ZMYM5</td><td>fw</td><td>AGAGTTGACTGAACAGACTCCT</td><td rowspan="2">PrimerBank ID: 218083691c1</td></tr>
<tr><td>rev</td><td>GACCAAATGAATCCCCTATGTCC</td></tr>
<tr><td rowspan="2">GPC5</td><td>fw</td><td>GGTGTGACTGACAGTTCCCTG</td><td rowspan="2">PrimerBank, ID: 215272348c3</td></tr>
<tr><td>rev</td><td>TGCAGATAGTCTGTGGTGTTGAT</td></tr>
<tr><td colspan="5">Control primer</td></tr>
<tr><td rowspan="2">ALB (chr4)</td><td>fw</td><td>TGTTGCATGAGAAAACGCCA</td><td rowspan="2" colspan="2">Bremer et al. 2015</td></tr>
<tr><td>rev</td><td>GTCGCCTGTTCAACCAAGGAT</td></tr>
</table>

**Supplementary Table 4.** QPCR primers used to identify whole and partial chromosome deletions. Primer binding sites for chr. 6p and 13q are illustrated in Extended Data Figure 7b.

| Name | Sequence | | Reference |
|---|---|---|---|
| **p53** | fw | CACCGACTTCCTGAAAACAACGTTC | Giacomelli et al. 2018 |
| | rev | AAACGAACGTTGTTTTCAGGAAGTC | |
| **p21** | fw | CACCGCCGCGACTGTGATGCGCTAA | McKinley and Cheeseman 2017 |
| | rev | AAACTTAGCGCATCACAGTCGCGGC | |
| **p31$^{comet}$** | fw | CACCGACTTGAGACAAGCTCTACGC | Thu et al. 2018 |
| | rev | AAACGCGTAGAGCTTGTCTCAAGTC | |
| **CDC16** | fw | CACCGCTCTAGATAACCGAACCC | This paper |
| | rev | AAACGGGTTCGGTTATCTAGAGC | |

**Supplementary Table 5.** SgRNAs used in this study for the generation of heterozygous and homozygous knockout cell lines. Underlined sequence represents added overhangs for the creation of dsDNA oligos that can be cloned into the gRNA/Cas9 plasmid, see Methods.

| Name | Sequence | |
|---|---|---|
| **p53** | fw | TTATAGGGAGGTCAAATAAGCAGCA |
| | rev | ATCTACAAGCAGTCACAGCACAT |
| **p21** | fw | GCCCGGCCAGGTAACATAGTG |
| | rev | GTGACAGGTCCACATGGTCTTC |
| **p31$^{comet}$** | fw | GCGTATGTCGGAGTGCCTGC |
| | rev | GTGCTTAAGCTGTTCATAGG |
| **CDC16** | fw | CTATGATCGCACCACTGAACTC |
| | rev | TGTCAGCATGTGATGTGATGTT |

**Supplementary Table 6.** Genotyping primers used for the identification and analysis of generated knockout cell lines.

| Name | Sequence |
|---|---|
| **Scrambled** | Scramble siRNA (Dharmacon/smartpool format - D-001810-10-05) |
| **MAD2** | siMAD2 (Dharmacon/smartpool format – L-003271-00-0005) |

**Supplementary Table 7.** SiRNA used in this study for the depletion of MAD2.
